# Modular genetic control of social status in a cichlid fish

**DOI:** 10.1101/2020.04.03.024190

**Authors:** Beau A. Alward, Vibhav Laud, Christopher J. Skalnik, Ryan A. York, Scott Juntti, Russell D. Fernald

## Abstract

Social hierarchies are ubiquitous in social species and profoundly influence physiology and behavior. Androgens like testosterone have been strongly linked to social status, yet the molecular mechanisms regulating social status are not known. The African cichlid fish *Astatotilapia burtoni* is a powerful model species for elucidating the role of androgens in social status given their rich social hierarchy and genetic tractability. Dominant *A. burtoni* males possess large testes, bright coloration, and perform aggressive and reproductive behaviors while non-dominant males do not. Social status in *A. burtoni* is in flux, however, as males alter their status depending on the social environment. Due to a teleost-specific whole-genome duplication, *A. burtoni* possess two androgen receptor (AR) paralogs, *ARα* and *ARβ*, providing a unique opportunity to disentangle the role of gene duplication in the evolution of social systems. Here, we used CRISPR/Cas9 gene editing to generate AR mutant *A. burtoni* and performed a suite of experiments to interrogate the mechanistic basis of social dominance. We find that *ARβ*, but not *ARα*, is required for testes growth and bright coloration, while *ARα*, but not *ARβ*, is required for the performance of reproductive behavior and aggressive displays. Both receptors are required to reduce flees from females and either AR is sufficient for attacking males. Thus, social status in *A. burtoni* is inordinately dissociable and under the modular control of two AR paralogs. This type of non-redundancy may be important in facilitating social plasticity in *A. burtoni* and other species whose social status relies on social experience.

**Significance Statement:** Social rank along a hierarchy determines physiological state and behavioral performance. A ubiquitous feature of social hierarchies is the communication of rank through non-physical signaling systems (e.g., coloration) and aggression, traits that correlate with the reproductive status of an individual. Despite the links identified between social status, physiology, and behavior, the molecular basis of social status is not known. Here, we genetically dissect social status in the African cichlid fish *Astatotilapia burtoni* using CRISPR/Cas9 gene editing. We show that two distinct androgen receptor (AR) genes control social status in a highly modular manner. This type of coordination of social status may be fundamental across species that rely on social information to optimally guide physiology and behavior.

## Introduction

Social animals often organize into hierarchies, wherein social status or rank determines many physiological and behavioral traits (1). Successful navigation through a social hierarchy requires the constant integration of social cues to optimize chances of survival and reproductive opportunities (2). Within these social hierarchies dominant and non-dominant individuals exist (1). Dominant individuals typically behave more aggressively and have more mating opportunities than non-dominant individuals. Dominant animals may also express conspicuous signals that show their status to others. This can take the form of bright coloration or body size. Dominant individuals have an activated reproductive system as indicated by their large gonads. On the other hand, non-dominant individuals are not aggressive and have very few, if any, chances to mate. Non-dominant individuals may appear inconspicuous to avoid confrontations from higher ranking individuals and possess small gonads. For some species, social hierarchies are in flux as non-dominant animals can ascend to dominant rank given the social opportunity (3). Despite what is known about the importance of social hierarchies in controlling traits that relate to reproduction, the molecular mechanisms underlying social status are unclear.

One candidate molecular substrate for regulating social status is the androgen signaling system. Dominant animals tend to have higher levels of androgens like testosterone compared to non-dominant animals. Pharmacological manipulations across species suggest that androgen signaling is required to enhance the motivation to seek higher social status (4–6). However, results of studies using pharmacology to tease apart the molecular mechanisms of social status are limited. For instance, pharmacological agents have off-target effects that are difficult if not impossible to control for (7). For this reason, genetic manipulations are typically preferred over pharmacological ones to dissect the molecular basis of complex physiological and behavioral functions. However, genetic models of social status have until recently not been available for species in which rich social hierarchies can be controlled in the laboratory.

The African cichlid fish *Astatotilapia burtoni* is a powerful model species for the genetic dissection of social status (3, 8, 8). In the laboratory, as in nature, male *A. burtoni* exist as either non-dominant or dominant, exhibiting clear variation in testes mass, coloration, and behavior (Fig. 1A-B). Pharmacological studies suggest AR signaling enhances reproductive but not aggressive behavior associated with social dominance in *A. burtoni* (5, 10). Additionally, due to a whole-genome duplication (WGD) event specific to teleost fish (11), *A. burtoni* possess two AR paralogs, *ARα* and *ARβ* (Table S1), providing a unique opportunity to disentangle the role of gene duplication in the evolution of social systems (12). Finally, recent work used CRISPR/Cas9 gene editing to generate mutant *A. burtoni*. With this in mind, we used gene editing to produce novel *A. burtoni* AR mutants and performed a suite of experiments interrogating the molecular basis of social dominance.

**Figure 1.**
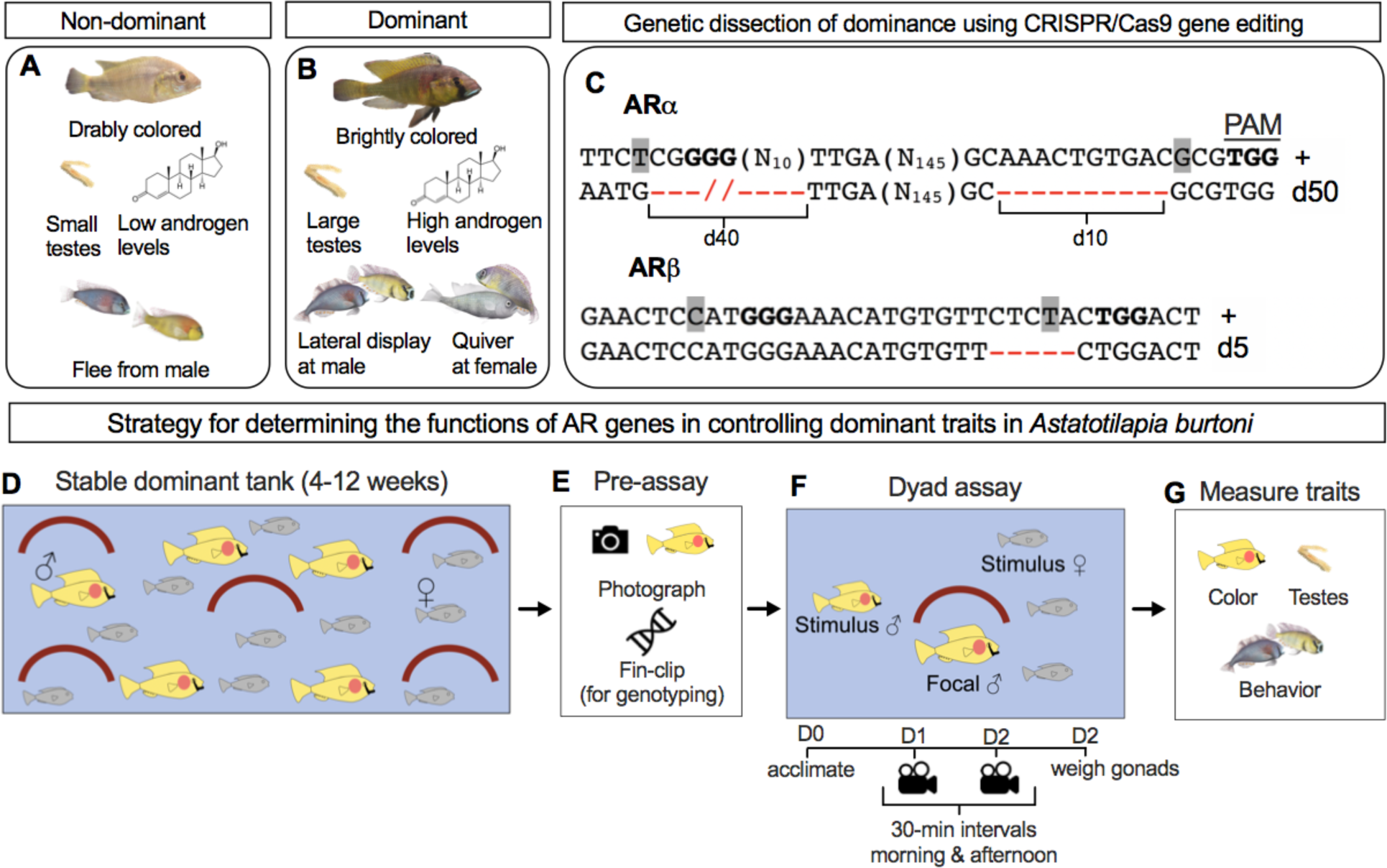
Understanding the control of social dominance by androgen receptors in *Astatotilapia burtoni*. **(A)** Non-dominant and **(B)** dominant male *A. burtoni* differ in terms of a variety of traits that reflect their social status. **(C)** We used CRISPR/Cas9 gene editing to generate *A. burtoni* that possess frameshift *ARα* or *ARβ* alleles. (**D** to **G**) Experimental strategy for testing the functions of *ARα* and *ARβ* in the control of social dominance. Predicted Cas9 cleavage sites are located to the left of letters highlighted in gray. Bolded 3-letter sequences indicate a PAM sequence. PAM=Protospacer Adjacent Motif. +=wild-type. d=deletion. (N_#_) indicate the number of bases not shown in actual gene sequence for clarity.

## Results

### Genetic dissection of social dominance in *A. burtoni* using CRISPR/Cas9 gene editing

We generated mutant fish possessing frameshift *ARα* or *ARβ* alleles using CRISPR/Cas9 gene editing (13). Two single-guide RNAs against *ARα* or *ARβ* and Cas9 mRNA (14, 15) were injected into embryos at the single-cell stage (16) after which frameshift mutant alleles were identified for both genes. *ARα* mutants possessed a total deletion of 50 basepairs (bp) (*AR*α^*Δ*^) and *ARβ* mutants possessed a deletion of 5 bp (*ARβ*^*Δ*^) (Fig. 1C). For the wild-type and mutant alleles we determined the predicted amino acid sequences and protein tertiary structure (Fig. S1) (17). Both mutant alleles contained premature stop codons that yielded predicted truncated amino acid sequences compared to wild-types (Fig. S1A-B). Analysis of the predicted tertiary structure of wild-type and mutant AR*α* and AR*β* revealed profound differences, with mutant versions of each protein totally lacking the complex tertiary structure observed in the wild-type versions (Fig. S1C-D). Thus, the frameshift alleles we generated for *AR*α and *ARβ* are highly likely to be completely non-functional. G_0_ injected fish were outcrossed with wild-type fish to generate heterozygous mutants (that were subsequently intercrossed). Heterozygous mutants of either genotype (*ARα*^*Δ/+*^ or *ARβ*^*Δ/+*^) were crossed to yield offspring of homozygous wild-types (*ARα*^*+/+*^ or *ARβ*^*+/+*^), heterozygous mutants (*ARα*^*Δ/+*^ or *ARβ*^*Δ/+*^), and homozygous mutants (*ARα*^*Δ/Δ*^ or *ARβ*^*Δ/Δ*^).

Do AR mutants display altered patterns of social dominance? To address this, we housed adult male fish from each cross for 4-12 weeks in stable dominant tanks (Fig. 1D), a housing environment that has been shown previously to reliably permit the full suite of social dominance traits (5, 18, 18). Fish were then assayed for three key traits related to social dominance: testes mass, coloration, and behavior (3) (Fig. 1E-G). First, individuals were removed and photographed (Fig. 1E) followed by fin-clip removal (for subsequent genotyping). Each fish was then placed into a dyad assay tank (Fig. 1E-F). In a dyad assay a focal fish (in this case, a fish from the stable dominant tank) is housed with a smaller stimulus male, three females, and a terra cotta pot simulating a potential mating site (20) (Fig. 1F). For the next two days fish were recorded from 8am-2pm. At 2 pm on the second day, focal fish were removed from the dyad assay tank and standard length, body mass, and testes mass were recorded. Blood was also collected for analysis of androgen levels. There was no effect of *AR*α or *ARβ* genotype on standard length or body mass (Fig. S2).

### *ARα* and *ARβ* are necessary for distinct aspects of social dominance

Variation in testes mass was measured first. We found that *ARα* and *ARβ* are required for normal testes mass, but in different directions. For instance, *ARα*^*Δ/Δ*^ males had testes that were larger than those seen in *ARα*^*+/+*^ males (Fig. 2A). On the other hand, *ARβ*^*Δ/Δ*^ and *ARβ*^*Δ/+*^ males had smaller testes than *ARβ*^*+/+*^ males and *ARβ*^*Δ/+*^ males had larger testes than *ARβ*^*Δ/Δ*^ males (Fig. 2B). Based on these findings, we predicted that *ARβ* mutants would lack dominant-typical coloration (16).

**Figure 2.**
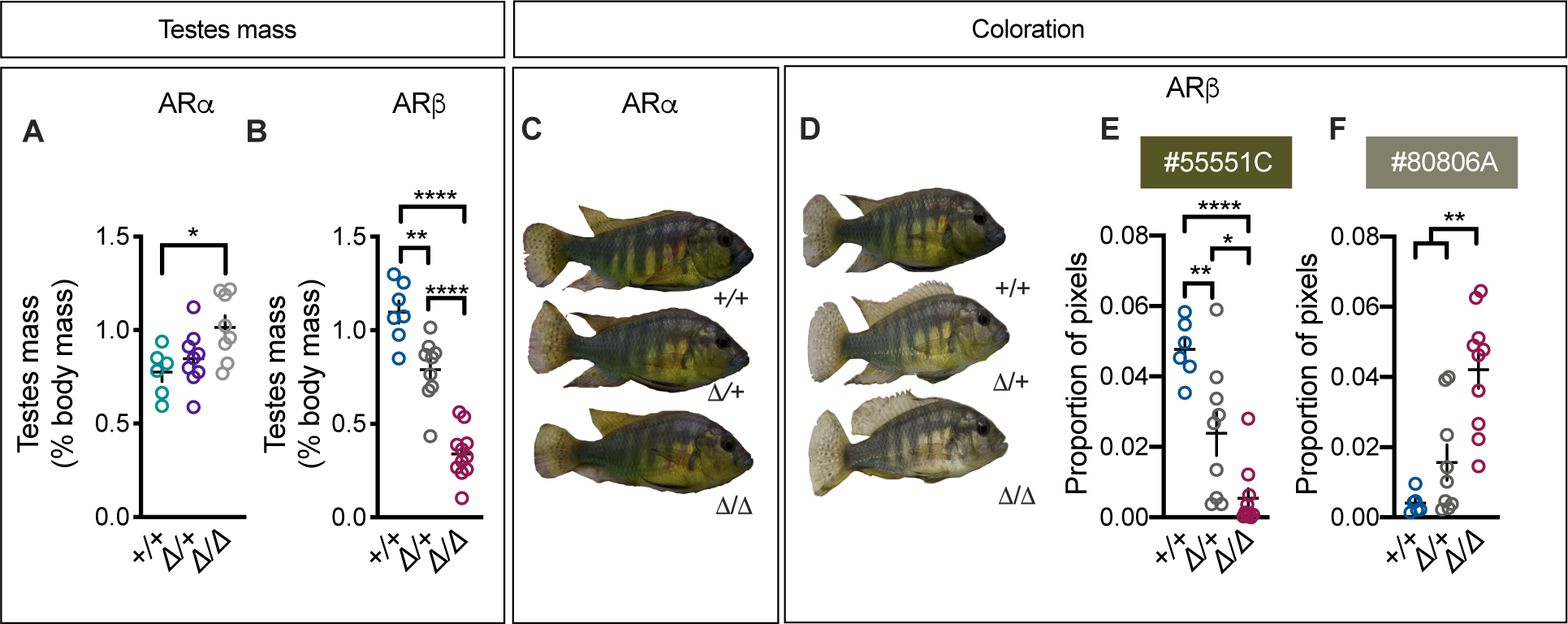
AR*α* and AR*β* play distinct roles in the control of testes mass and coloration. **(A)** *AR*α^*+/+*^ males have smaller testes than *AR*α^*Δ/Δ*^ males, while *AR*α^*Δ/+*^ males were not different than either group. **(B)** *ARβ*^*+/+*^ males have larger testes than *ARβ*^*Δ/+*^ males, which have larger testes than *ARβ*^*Δ/Δ*^ males. **(C)** *ARα* mutant males do not look different than *AR*α^*+/+*^ males (D) while *ARβ* mutant males lack the dominant-typical yellow coloration seen in *ARβ*^*+/+*^ males. **(E)** *ARβ*^*+/+*^ males possessed more “very dark yellow” (#55551C) pixels than *ARβ*^*Δ/+*^ males, which had more dark yellow pixels than *ARβ*^*Δ/Δ*^ males, **(F)** while *ARβ*^*Δ/Δ*^ males had more “dark grayish yellow” (#80806A) pixels than both *ARβ*^*Δ/+*^ and *ARβ*^*Δ/Δ*^ males. +=wild-type. *Δ*=frameshift deletion. Circles represent data points for individual fish. Crosses represent Mean±SEM. \*\*\*\**P*<0.0001; \*\**P* < 0.01; \**P* < 0.05.

Dominant male *A. burtoni* typically express bright yellow or blue coloration while non-dominant males appear drab (3) (Fig. 1A-B). Visual inspection showed that, while *ARα*^*Δ/+*^ and *ARα*^*Δ/Δ*^ males looked no different than *ARα*^*+/+*^ males (Fig. 2C), *ARβ* mutant males looked profoundly different than *ARβ*^*+/+*^ males, which possessed dominant-typical yellow coloration (3) (Fig. 2D). Hierarchical clustering confirmed these observations (Fig. S3 and Fig. S4). To determine what specific colors were different between *ARβ* mutant males and *ARβ*^*+/+*^ males, we performed quantitative analysis on images of each fish. *ARβ*^*+/+*^ males differed significantly from *ARβ* mutant males for several colors (Fig. 2E-F; Fig. S5). For example, *ARβ*^*+/+*^ males possessed more “very dark yellow” (hex: #55551C) pixels than *ARβ*^*Δ/+*^ males, which had more dark yellow pixels than *ARβ*^*Δ/Δ*^ males (Fig. 2E), while *ARβ*^*Δ/Δ*^ males had more “dark grayish yellow” (#80806A) pixels than both *ARβ*^*Δ/+*^ and *ARβ*^*Δ/Δ*^ males (Fig. 2F). Therefore, *ARβ* is required for dominant coloration, while *ARα* is not.

Given the above observations we formed several hypotheses for what to expect from the behavior analysis of mutant fish. Specifically, we anticipated that *ARα*^*Δ/Δ*^ and *ARα*^*Δ/+*^ males should exhibit normal levels of dominant behavior, while *ARβ*^*Δ/+*^ and *ARβ*^*Δ/Δ*^ fish should exhibit decreased levels of dominant behavior (3). Strikingly, the exact opposite was true. All reproductive behaviors in *ARα* mutant males were virtually abolished (Fig. 3A; Fig. S6A-D). *ARα* mutant males performed significantly fewer aggressive displays than *ARα*^*+/+*^ males (Fig. 3B), while attacks directed towards males were unaffected in *ARα* mutant males (Fig. S6E-F). *ARα* mutant males did not differ from *ARα*^*+/+*^ males in the number of times they fled from the stimulus male (Fig. S6G); however, *ARα*^*Δ/Δ*^ males fled significantly more from females compared to *ARα*^*+/+*^ males (Fig. 3C). Importantly, there was no effect of *ARα* genotype on how often the stimulus females bit the focal male (Fig. 3D), suggesting the effects on fleeing from females were not due to higher aggression from the stimulus female towards the AR mutant focal males. *ARβ* mutant fish did not differ from *ARβ*^*+/+*^ males for all behaviors (Fig. 3E-F; Fig. S6) except for one, flee from female. Specifically, *ARβ* mutants fled from females more than *ARβ*^*+/+*^ males (Fig. 3G), coinciding with the findings for this behavior in *ARα* mutants (Fig. 3C). As with *ARα*, there was no effect of *ARβ* on how often the stimulus females bit the focal male (Fig. 3H). Thus, both *ARα* and *ARβ* are required to reduce flees from females.

**Figure 3.**
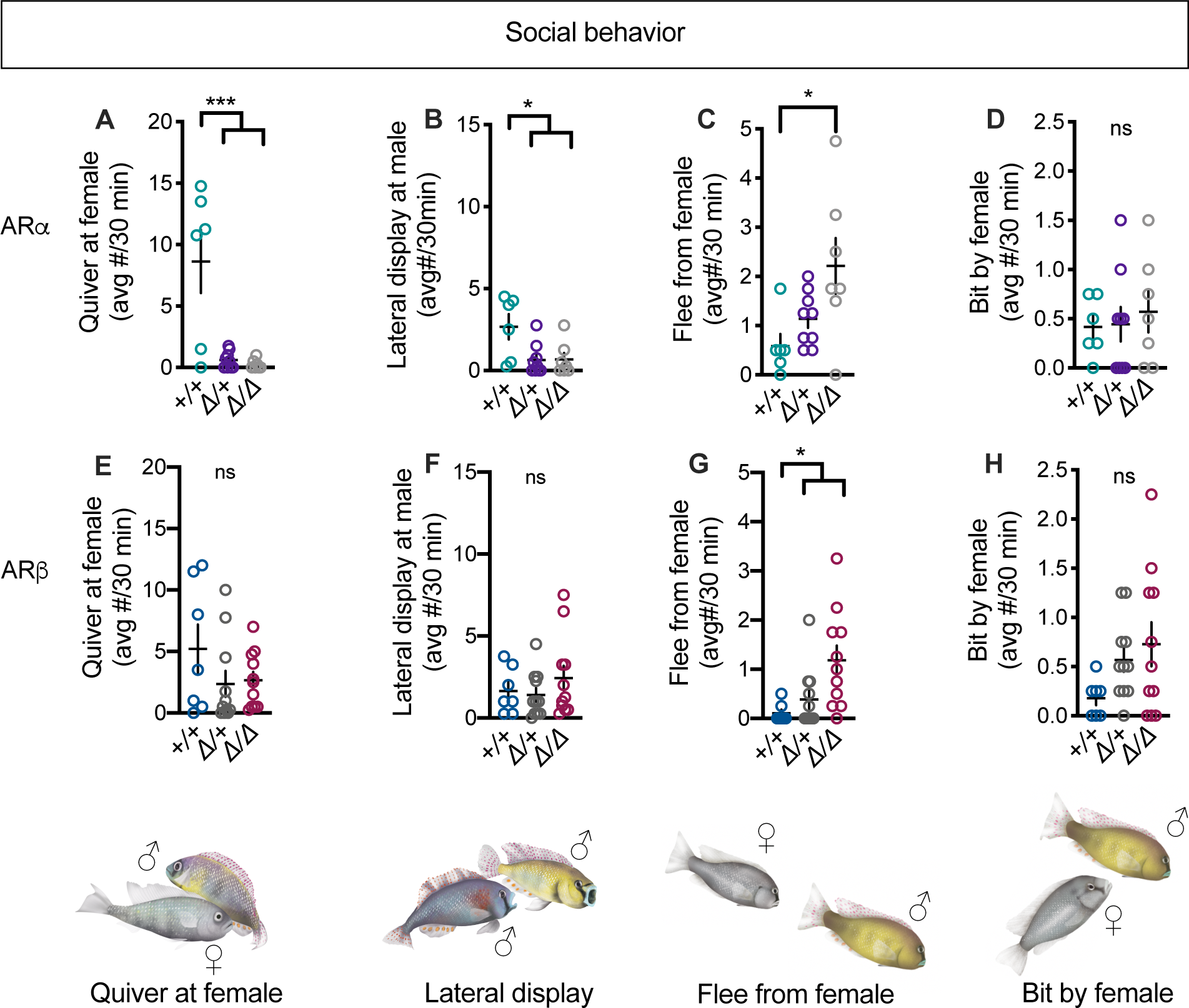
Dissociable roles of AR*α* and AR*β* in the regulation of social behavior. **(A)** *AR*α mutant males quivered at females significantly less than *AR*α^*+/+*^ males. **(B)** *AR*α mutant males performed lateral displays at males significantly less than *AR*α^*+/+*^ males. **(C)** *AR*α^*Δ/Δ*^ males fled from females significantly more than *AR*α^*+/+*^ males, **(D)** but there was no effect of *AR*α genotype on the number of times focal males were bit by females. (**E** and **F**) There was no effect of *ARβ* genotype on quiver at female or lateral display at female, **(G)** but *ARβ* mutants fled from females more than *ARβ*^*+/+*^ males. (H) There was no effect of *ARβ* genotype on the number of times focal males were bit by females. +=wild-type. *Δ*=frameshift deletion. Circles represent data points for individual fish. Crosses represent Mean±SEM. \*\*\**P* < 0.001; \**P* < 0.05.

### Analysis of AR double mutants reveal either *ARα* or *ARβ* is sufficient for attacking males

We were surprised that *ARα* was required for the performance of an aggressive display, but neither it nor *ARβ* was required for attacking other males. This suggests 1) either AR is sufficient for attacking other males, or 2) an AR-independent mechanism controls attacks directed towards males. We were able to test the first hypothesis by generating AR double mutants through breeding strategies. Given that some *ARα* mutant males were severely injured in the dyad assay (see Supplementary Results), we used a modified dyad assay in which the focal fish was isolated physically from the three females and stimulus male using two transparent, perforated plastic barriers on either side (Fig. S7A). This approach is sufficient for testing our hypothesis about the role of either AR in controlling aggression specifically given previous work (5, 18) showing males perform aggressive displays at males on the other side of a barrier and also attack the stimulus male through the barrier (Fig. S7B-C).

We assayed four AR double wild-type (*ARα*^*+/+*^*;ARβ*^*+/+*^) males and six AR double mutant (*ARα* ^*Δ/Δ*^*;ARβ*^*Δ/Δ*^) fish. Three of the *ARα*^*Δ/Δ*^*;ARβ*^*Δ/Δ*^ fish were found to be female, which was not determined until dissection where the presence of ovaries was confirmed. *ARα*^*+/+*^*;ARβ*^*+/+*^ males weighed more than *ARα* ^*Δ/Δ*^*;ARβ*^*Δ/Δ*^ males and females (Fig. S8A) and were greater in standard length than *ARα* ^*Δ/Δ*^*;ARβ*^*Δ/Δ*^ males (Fig. S8B), suggesting that AR regulates body size in line with findings in AR mutant mice (*18*). Like *ARβ* mutant males, *ARα* ^*Δ/Δ*^*;ARβ*^*Δ/Δ*^ males had extremely small testes compared to *ARα*^*+/+*^*;ARβ*^*+/+*^ males (Fig. 4A). As with *ARβ* mutants, *ARα* ^*Δ/Δ*^*;ARβ*^*Δ/Δ*^ males lacked the dominant-typical coloration seen in *ARα*^*+/+*^*;ARβ*^*+/+*^ males and were indistinguishable from *ARα* ^*Δ/Δ*^*;ARβ*^*Δ/Δ*^ females (Fig. 4B). Hierarchical clustering confirmed this observation (Fig. S9). Quantitative image analysis revealed that *ARα*^*+/+*^*;ARβ*^*+/+*^ males differed from *ARα* ^*Δ/Δ*^*;ARβ*^*Δ/Δ*^ males and females for several colors (Fig. 4C-D; Fig. S10). For example, *ARα*^*+/+*^*;ARβ*^*+/+*^ males possessed more “very dark grayish lime green” (#475547) and “dark grayish lime green” (#6A806A) pixels than *ARα* ^*Δ/Δ*^*;ARβ*^*Δ/Δ*^ males and females, which were indistinguishable from one another (Fig. 4C-D).

**Figure 4.**
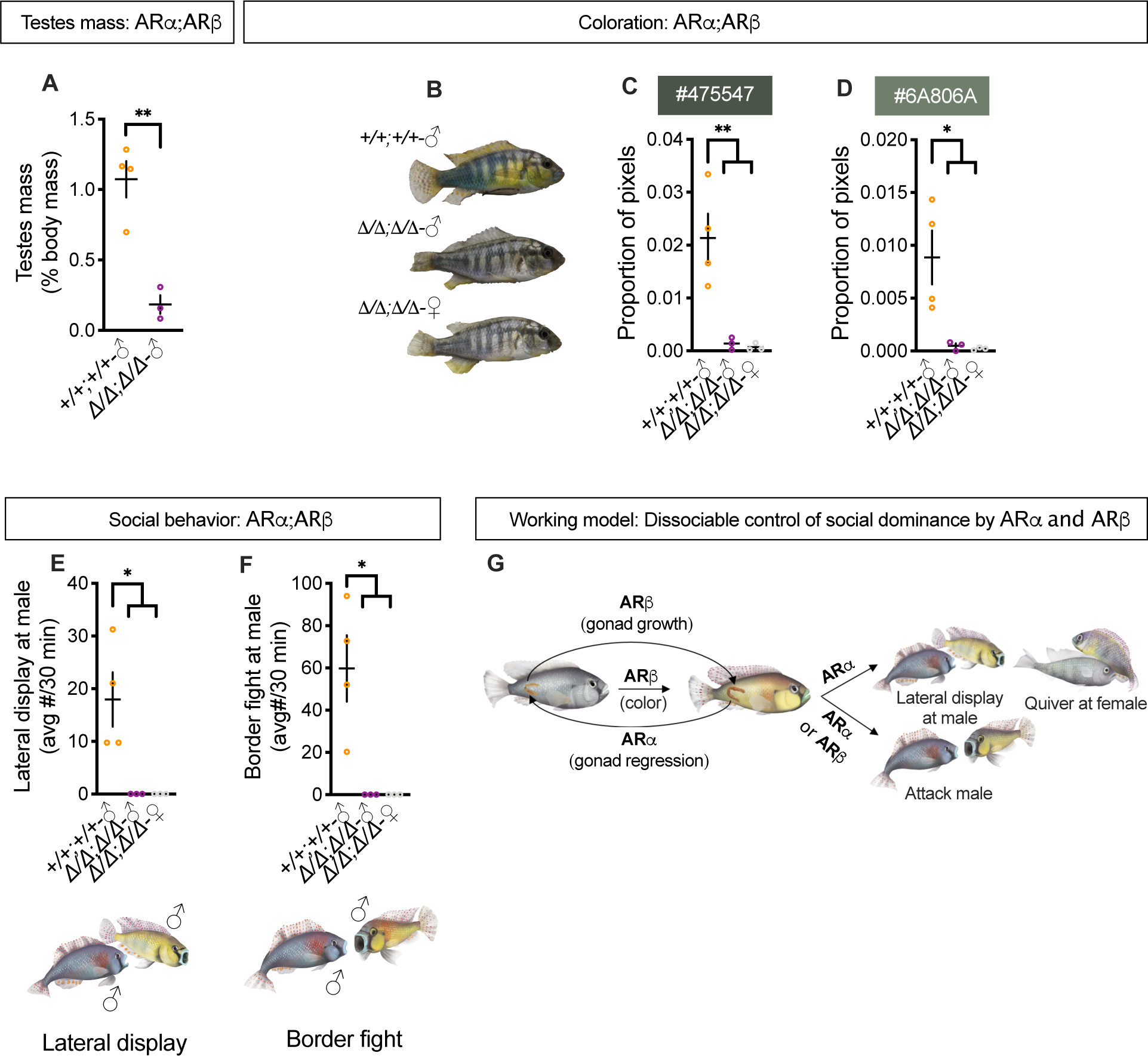
Analysis of AR double mutants highlights the modular control of social dominance by *ARα* and *ARβ*. **(A)** *ARα* ^*Δ/Δ*^*;ARβ*^*Δ/Δ*^ males had smaller testes than *ARα*^*+/+*^*;ARβ*^*+/+*^ males. **(B)** *ARα* ^*Δ/Δ*^*;ARβ*^*Δ/Δ*^ males and females lacked the dominant-typical coloration observed in *ARα*^*+/+*^*;ARβ*^*+/+*^ males. **(C)** *ARα*^*+/+*^*;ARβ*^*+/+*^ males possessed more “very dark grayish lime green” (#475547) **(D)** and “dark grayish lime green” (#6A806A) pixels than *ARα* ^*Δ/Δ*^*;ARβ*^*Δ/Δ*^ males and females, which were indistinguishable from one another. **(E)** *ARα*^*+/+*^*;ARβ*^*+/+*^ males performed more lateral displays **(F)** and border fights directed towards the stimulus male than *ARα* ^*Δ/Δ*^*;ARβ*^*Δ/Δ*^ males and females. **(G)** A summary of the current findings in the form of a working model. +=wild-type. *Δ*=frameshift deletion. Circles represent data points for individual fish. Crosses represent Mean±SEM. \*\**P* < 0.01; \**P* < 0.05.

In addition to the effects described above, we observed striking differences in aggressive behavior between *ARα*^*+/+*^*;ARβ*^*+/+*^ and *ARα* ^*Δ/Δ*^*;ARβ*^*Δ/Δ*^ fish (Fig. 3E-H). *ARα*^*+/+*^*;ARβ*^*+/+*^ males performed significantly higher levels of aggressive displays and attacks directed towards the stimulus male compared to *ARα* ^*Δ/Δ*^*;ARβ*^*Δ/Δ*^ males and females (Fig. 4H-I). Indeed, none of the *ARα* ^*Δ/Δ*^*;ARβ*^*Δ/Δ*^ fish performed these behaviors. Surprisingly, both *ARα* ^*Δ/Δ*^*;ARβ*^*Δ/Δ*^ males and females performed lateral displays directed towards *females*, while none of the *ARα*^*+/+*^*;ARβ*^*+/+*^ males performed this behavior (Fig. S11A). *ARα* ^*Δ/Δ*^*;ARβ*^*Δ/Δ*^ males also directed attacks towards females more than *ARα*^*+/+*^*;ARβ*^*+/+*^ males (Fig. S11B), but this difference did not reach statistical significance (*P*=0.06). Previous work has shown that female *A. burtoni* perform these acts of aggression towards one another (21). Indeed, the stimulus females in our assays were seen attacking and performing lateral displays at one another (data not shown). Therefore, the aggressive behaviors performed by *ARα* ^*Δ/Δ*^*;ARβ*^*Δ/Δ*^ males and females appear to be female-typical. Our findings support our first hypothesis that either AR is sufficient for male-directed attacks, demonstrating further evidence for highly modular control of social dominance traits by AR genes.

### Observed effects on social dominance are not due to the absence of circulating androgens

To determine whether observed effects on dominance traits in AR mutant *A. burtoni* could have been be due to abnormally low levels of androgens (22, 23), we measured levels of testosterone and 11-ketotestosterone (11-KT) in all fish. *ARα*^*Δ/Δ*^ and *ARα*^*Δ/+*^ males had levels of testosterone and 11-KT that were indistinguishable from *ARα*^+*/+*^ males (Fig. S12A-B). *ARα*^*Δ/Δ*^ and *ARα*^*Δ/+*^ males had levels of testosterone that were not different from *ARα*^*+/+*^ males (Fig. S12C). *ARβ*^*Δ/Δ*^ males possessed higher levels of 11-KT compared to *ARβ*^*Δ/+*^ and *ARβ*^*+/+*^ males (Fig. S12D), which is in line with previous work in AR mutant female mice that have intact gonads and significantly higher levels of 5-alpha dihydrotestosterone, the functionally similar metabolite of testosterone found in mammals, compared to wild-types (24).

*ARα* ^*Δ/Δ*^*;ARβ*^*Δ/Δ*^ males and females had levels of testosterone that were not different from *ARα*^*+/+*^*;ARβ*^*+/+*^ males (Fig. S12E). Like *ARβ*^*Δ/Δ*^ males, *ARα* ^*Δ/Δ*^*;ARβ*^*Δ/Δ*^ males had significantly higher levels of 11-KT compared to *ARα*^*+/+*^*;ARβ*^*+/+*^ males, while *ARα* ^*Δ/Δ*^*;ARβ*^*Δ/Δ*^ females were not different from either group (Fig. S12F). These findings collectively indicate that the observed perturbations in dominance traits in AR mutants are not due to abnormally low levels of androgens.

## Discussion

These data present coherent evidence for remarkable modularity in the control of social status in male *A. burtoni*. We show a stark double dissociation in the regulation of social status, wherein *ARα* is required for most dominant behaviors and testes regression but not dominant coloration, while *ARβ* is required for dominant coloration and testes growth, but not dominant behaviors. At the same time, both *ARα* and *ARβ* are required for reducing flees from females and either AR is sufficient for males to attack other males (results summarized in Fig. 4J). Our findings have important implications for understanding social status and social behavior across species.

Like other species, male *A. burtoni* constantly survey the social environment waiting for an opportunity to achieve social dominance (2). Previous work in *A. burtoni* has shown males can uncouple specific dominance traits as a function of variation in social information (2, 3). For instance, non-dominant males will turn on bright colors and perform dominant behaviors in the absence of large testes when larger dominant males cannot see them, but immediately turn off their colors and cease dominant behaviors when the dominant male can see them (25). Dominant males are also able to uncouple coloration and behavior from physiological state in order to deceive neighboring males that are larger than them (26). The ability to dissociate coloration, physiology, and behavior depending on the social environment reflects a sophisticated social calculus that optimizes chances of survival and reproduction (2). Our results suggest that this social calculus may be controlled by distinct AR genes. Moreover, the current findings provide further support for theories stating that androgen signaling mediates complex suites of physiological and behavioral responses that contribute to successful social interactions (5, 6, 27, 28).

Through genetic dissection using CRISPR/Cas9 gene editing, we provide compelling evidence that *ARα* and *ARβ* have been subfunctionalized, the process by which each duplicated gene retains a subset of ancestral functions (12). For example, in zebrafish (*Danio rerio*), which possess a single AR gene, mutation of AR produces male fish that possess small testes, reduced color, and perform fewer courtship behaviors compared to wild-types (29, 30). The results in zebrafish parallel those found in AR mutant mice, which also possess a single AR gene (22). In *A. burtoni*, we have shown that the control of these classes of traits is distributed over *AR* paralogs. An intriguing follow-up question to our findings is whether the subfunctionalization of AR genes across different cichlid fish contributes to the rich array of social dynamics seen in this clade (11, 31). Given the ascent of genome editing tools like CRISPR/Cas9 gene editing, this question can now be tested in species with a range of different social systems and evolutionary trajectories to discover the fundamental molecular mechanisms of social status.

Based on our findings, we propose a working framework where the type of dissociable control of social status observed here occurs in other species that rely on social and environmental information to optimize physiology and behavior. In this framework, independent mechanisms integrate social cues and regulate distinct aspects of traits that relate to reproduction. This regulation may begin with androgen signaling, which acts on separate molecular and neural pathways that govern distinct dominance traits in a non-overlapping manner. In this way, our framework is fundamentally similar to the hormonal control of courtship birdsong, wherein discrete steroid-sensitive molecular and neural pathways that regulate different features of song are activated by androgen signaling during the breeding season (27). This framework can be usefully applied to studies aiming to understand how non-dominant and dominant animals rapidly alter social behavior in way that is seemingly disconnected from their reproductive state and vice-versa (1, 2). By performing rich mechanistic studies in a genetically-tractable model species of social status, we have yielded fundamental insights into the nature of social behavior.

## Materials and Methods

### Generation of frameshift alleles for *ARα* and *ARβ* using CRISPR/Cas9 gene editing

Fish were bred and used at Stanford University from a colony derived from Lake Tanganyika (32) in accordance with AAALAC standards.

We generated fish with mutant *ARα* or *ARβ* alleles using an approach similar to that of Juntti et al (16). Mutations of either gene were induced by injection of two single-guide RNAs (sgRNA) simultaneously targeting regions upstream from the DNA binding domain (DBD) and ligand-binding domain (LBD) within exon 1 for *ARα* and exon 2 for *ARβ*. For *ARα*, we designed sgRNAs targeting sequence *ARα-*A, 5’-ACTGTGGCGGATACTTCTCG-3’, and sequence *ARα-*B, 5’-GGTGCGCAAACTGTGACGCG-3’, whose cut sites were separated by 178 basepairs (bp). For *ARβ*, we designed sgRNAs targeting sequence *ARβ-*A, 5’-GGGAAACATGTGTTCTCTAC-3’, and *ARβ -*B, 5’-GGGGGAAAGAAGAACTCCAT-3’ whose cut sites were separated by 21 bp. We generated each sgRNA using cloning-free sgRNA synthesis (33). For instance, to synthesize sgRNA targeting *ARα-*A (g*ARα*-A) we annealed oligonucleotide-*ARα-*A (oligo-*ARα-*A), which contained the *ARα-*A target sequence, and oligo-2, a generic oligo that we used for all sgRNA synthesis reactions (see Table S2 for all oligo sequences). g*ARα*-B, g*ARβ*-A, and g*ARβ*-B were synthesized in the same manner.

We waited for 30 min of fertilization and then injected single-cell embryos with the two sgRNA targeting *ARα* or *ARβ*. We delivered ∼1 nl of each sgRNA, 60 ng/ml *nls-zCas9-nls* mRNA, and 0.3% Texas-Red-conjugated dextran (3,000 MW; Life Technologies). In ∼5-week embryos injected with g*ARα*-A and g*ARα*-B, we PCR amplified a 536-bp amplicon spanning the *ARα-*A and *ARα-*B target sites with the primers AR*α*FlankF, 5’-CCCAGTGCACTCTAACTCCG -3’ and AR*α*FlankR, 5’ - TTTAAGGGTACGACCTCGGC -3’, and Sanger sequenced the product with AR*α*FlankR (MCLabs). We performed the same procedure for embryos injected with g*ARβ*-A and g*ARβ*-B by PCR amplifying a 642-bp amplicon with the primers ARbFlankF, 5’ - CCATCCCACCTCCAAGAGTC -3’ and AR*β*FlankR, 5’ - GAGGACAGGCCGATGATGAA -3’, and Sanger sequenced the product with AR*β*FlankF (MCLabs). We saved fish showing evidence of mutations in *ARα* or *ARβ* and crossed these fish to wild-types. These G1 offspring carried a variety of indel alleles, so we selectively propagated an allele for each gene predicted to result in a loss of function (i.e., a 50-bp deletion for *ARα* and a 5-bp deletion for *ARβ*). We intercrossed G1 fish from different founders to obtain biallelic *ARα* mutants (*ARα*^*Δ*/*Δ*^), heterozygous *ARα* mutants (*ARα*^*Δ*/+^), and *ARα* wild-types (*ARα*^+/+^) or biallelic *ARβ* mutants (*ARβ* ^*Δ*/*Δ*^), heterozygous *ARβ* mutants (*ARβ* ^*Δ*/+^), and *ARβ* wild-types (*ARβ* ^+/+^).

### Establishing stable dominant tanks

Social dominance is reliably induced when males have ample opportunity to establish a territory and access to females to mate with (5, 34). This social opportunity can be established using stable dominant tanks, wherein 5-8 size-matched males are housed in a 32-gallon tank with 10-15 females and 5 potential mating sites that are represented by halved terra cotta pots. We housed males from either cross in separate stable dominant tanks for 4-12 weeks. Each stable dominant tank contained males that were matched by the particular cross they arose from. The age of males included in the dyad assays ranged from 6-10 months. 2-3 age- and size-matched males from a given stable dominant tank were ran through separate dyad assays simultaneously.

### Photography

Fish were removed from their tank and placed for ∼20 seconds on a white paper towel to dry them off. Then, fish were immediately placed onto a white paper towel within a light chamber that contained a ruler for scale and photographed using a Sony camera (Sony Alpha NEX-C3 16 MP; shutter speed=1/80; aperture=4.5; white balance=+0.0) mounted on a tripod. 6-10 photos were taken to increase the likelihood of capturing an image during which the fish were not operculating. Fish were fin-clipped (see “Genotyping”) and immediately moved to their dyad assay tank. This whole process took ∼45 seconds. Images were transferred from the camera SD card to a computer (Mac). We were unable to take photos of one *ARβ* ^+/+^ fish, two *ARβ* ^*Δ*/+^fish, and one *ARβ* ^*Δ*/*Δ*^ fish.

### Quantitative image analysis

JPEG images with the highest resolution and where the fish was not operculating were chosen for analyses. To perform quantitative analysis of photos, we first cropped the fish out of the rest of the image using the Lasso Selection tool in Preview. Cropped images were then analyzed using R code (colordistance package: https://CRAN.R-project.org/package=colordistance), which computes the proportion of pixels in an image occupied by each color. For *ARα* fish, if 15 or more fish (two-thirds of fish analyzed) had a zero value for a given color, this color was not considered for further analysis. For *ARβ* fish, if 16 or more fish had a zero value for a given color, this color was not considered for further analysis. For *ARα*;*ARβ* fish, if 7 or more fish had a zero value for a given color, this color was not considered for further analysis. We measured the proportion of pixels occupied by 84 colors for *ARα* fish, 83 colors for *ARβ* fish, and 76 colors for *ARα*;*ARβ* fish.

### Fin-clipping and DNA extraction

After photographing the fish, they were immediately fin-clipped. Using ethanol-cleaned scissors, a 1-2 mm portion of the anal fin was excised and placed into an individual PCR tube. This was repeated for the rest of the fish ran on a given day and the scissors were cleaned thoroughly with ethanol between each fin-clipping. To extract DNA, 180 ml of NaOH (50 mM) was added to the sample, which was incubated at 94 C for 15 minutes. Then, 20 ml of Tris-HCL (ph=8) was added directly into the sample, which was then vortexed and spun down using a minicentrifuge for 5 seconds. The samples were then placed at -20C for at least 15 minutes before PCR amplification of mutated regions of *ARα* or *ARβ* (see “Generation of frameshift alleles for ARa and ARb using CRISPR/Cas9 gene editing”).

### Dyad assay setup

Each dyad assay was conducted in 8-gallon tanks with enough gravel spread evenly to cover the bottom of the tank and a half terra cotta pot (simulating a potential mating site) in the middle. An air stone supplying oxygen to the water was present in each tank. Each dyad assay tank was visually isolated from nearby tanks using black plastic barriers between and behind tanks. After fish were photographed and fin-clipped (see Fin-clipping and DNA extraction) they were immediately transferred to the tank. Three stimulus females and one stimulus male were collected from community tanks and then added to the dyad assay tank. We aimed to always include gravid females but if this could not be accomplished 1-2 females who were obviously not brooding were included. Stimulus males were chosen based on being smaller than the focal male based on visual inspection. The standard length of the stimulus male was measured after each assay and ranged from being 8-25% smaller than the focal male. Notably, before the assay the stimulus females had never interacted visually, chemically, or physically with the stimulus or focal males and neither had the focal and stimulus male interacted with each other. This process was then repeated for all the other males for which dyad assays were ran. 2-3 dyad assays were ran simultaneously.

A Wifi-enabled camcorder (Canon VIXIA HF R80) was then mounted on tripod placed in front of each tank. Recording began the next day (day 1) at 9am and was started remotely using the Wifi function, preventing any disturbance from the experimenter to start recording. Recordings were stopped remotely at 2pm when the fish were fed. The day after (day 2), recording commenced in the same fashion except at 2pm focal fish were removed and tissue was harvested for physiological measurements and blood was collected.

### Scoring behavior

Behavior was scored during 30-minute intervals on day 1 and day 2. The first 30 minutes of scoring for day 1 and day 2 started from the first observation of behavior performed by the focal male after lights on. If an hour elapsed and the fish did not perform a behavior, scoring in the morning was stopped and zero occurrences was recorded for all behaviors for that time point. On day 1 and day 2, the final 30 minutes before lights on—from 1:30 to 2:00 pm—were also scored. Based on previous work, multiple types of behavior were quantified (32): subordinate behavior (flee from male or flee from female); territorial or agonistic behaviors (lateral display, chase male, bite male, and bite female); and reproductive behaviors (chase female, quiver, lead swim, pot entry, and dig). Fleeing was defined as a rapid retreat swim from an approaching fish. Lateral displays are aggressive displays classified as presentations of the side of the body to another fish with erect fins, flared opercula, and trembling of the body. Biting was defined as the male lunging a short distance towards a fish and biting it on its side and floating backwards a short distance. Chase was defined as a rapid swim directed towards a fish. Chase male, bite male, and bite female are considered attack behaviors. Chase female was grouped with reproductive behaviors because they are a normal component of the courtship repertoire. Quiver was defined as a rapid vibration of the body by the male with presentation of the anal fin egg spots to a female, and lead swim was defined as swimming towards the shelter accompanied by back-and-forth motions of the tail (waggles) as the male attempted to lead a female towards the pot. We defined pot entry as any time the focal male entered the half terra cotta pot and digging as any time the male scooped gravel from inside its pot or around its pot into its mouth and subsequently released it around its pot. Videos were scored in Scorevideo (Matlab). The results of scoring videos were saved into log files that were subjected to a variety of analyses using custom R software. Behaviors across the four 30-minute intervals were averaged (average # of behavior/30 minutes) and statistical analyses were performed on these values.

### Modified dyad assay setup and scoring

Fish included in the modified dyad assay setup were handled in the same way as in the normal dyad assay setup in terms of photography and fin-clipping. The tanks were setup differently, however. Two perforated, transparent, acrylic barriers were placed in the 8-gallon tank to separate the focal fish from three females on one side and one stimulus male on the other. We scored multiple aggressive behaviors in this setup. As with the normal dyad assay setup, we scored aggressive displays: lateral displays that directed at the stimulus male or the stimulus females. We also scored attack behaviors called border fights, which involved the focal fish attacking the stimulus fish across the acrylic barrier and is typified by head-on lunges and rams against the barrier with an open mouth.

### Morphological and steroid hormone analyses

Focal fish were assessed for standard length, body mass, and testes mass (corrected for body mass). Blood samples were also collected with capillary tubes from the caudal vein, centrifuged for 10 min at 5200g, and the plasma was removed and stored at –80 °C until assayed. Immediately after blood collection fish were killed by cervical transection. Testes were removed and weighed. Testes mass could not be recorded for one *AR*α^*Δ/+*^ and one *AR*α^*Δ/Δ*^ male that died before the end of the assay. SL, BM, and testes mass were not recorded for two *ARβ*^*Δ/+*^ males.

Plasma testosterone and 11-ketotestosterone (11-KT) levels were measured using commercially available enzyme immunoassay (EIA) kits (Cayman Chemical Company, Ann Arbor, MI, USA) as previously described and validated for this species (Maruska and Fernald, 2010b). Briefly, for testosterone and 11-KT assays, a 1-5 μl sample of plasma from each subject was extracted three times using 200 μl of ethyl ether and evaporated under a fume hood before re-constitution in EIA assay buffer. EIA kit protocols were then strictly followed, plates were read at 405 nm using a microplate reader (UVmax Microplate Reader; Molecular Devices, Sunnyvale, CA, USA) and steroid concentrations were determined based on standard curves. All samples were assayed in duplicate. Two *ARβ*^*Δ/Δ*^ males could not be assayed for testosterone or 11-KT due to possible contamination and blood was not collected for one *ARβ*^*Δ/+*^ male. One *AR*α^*Δ/+*^ and one *AR*α^*Δ/Δ*^ male could not be assayed for testosterone or 11-KT because they died before the end of the assay (see Main Text). One *AR*α^*+/+*^ and one *AR*α^*Δ/Δ*^ male could not be assayed for testosterone or 11-KT because their plasma was mistakenly discarded.

### Statistics and clustering

All statistical tests were performed in the R statistical computing environment or Prism 8.3. We used One-Way ANOVAs for all traits tested. Following a significant main effect for an ANOVA, Tukey’s post-hoc tests were used for pairwise comparisons. An individual t-test was used to compare testes mass between *ARα*^*+/+*^*;ARβ*^*+/+*^ males and *ARα*^*Δ/Δ*^*;ARβ*^*Δ/Δ*^ males. Differences were considered significant at p≤0.05.

To cluster fish based on their coloration, we computed the Euclidean distances between all animals using the R function *dist*. We input these distances into the R function *hclust*.

Complete linkages were used to build the hierarchical dendrograms in Fig. S3, Fig. S4, and Fig. S10.

## Acknowledgments

We thank Danielle Blakkan for assistance with the experimental set-up and Andrew Hoadley for assistance with hormone extractions and EIAs and providing feedback on an earlier version of this manuscript.

## Funding

This work was supported by an Arnold O. Beckman Fellowship to B.A.A., a University of Houston-National Research University Fund (NRUF: R0503962) to B.A.A., an NIH NS034950, NIH MH101373, and NIH MH 096220 to R.D.F., and S.A.J. is supported by an EDGE grant (NSF IOS-1825723).

## Supplementary Information

### Supplementary Results

#### Injuries to focal males

We found that 2 *AR*α^*Δ/wt*^ males and 1 *AR*α^*Δ/Δ*^ had damaged fins; 1 injured fish from each of these respective groups died on day 2 before the extractions. Injuries to these fish were likely caused by attacks from the stimulus male as these males fled at high levels from males compared to fish that were not injured (average flee from male±SEM= 74.4±43.29 and 12.16±5.72 for injured and uninjured fish, respectively) and fled from females at levels similar to uninjured fish (average flee from female±SEM= 0.83±0.60 and 1.41±0.26 for injured and uninjured fish, respectively. No *AR*α^*+/+*^ males exhibited injuries and all survived the assay. No injuries were observed in *ARβ*^*+/+*^, *ARβ*^*Δ/wt*^, or *ARβ*^*Δ/Δ*^ fish.

### Supplementary Figures and Legends

**Fig. S1.**
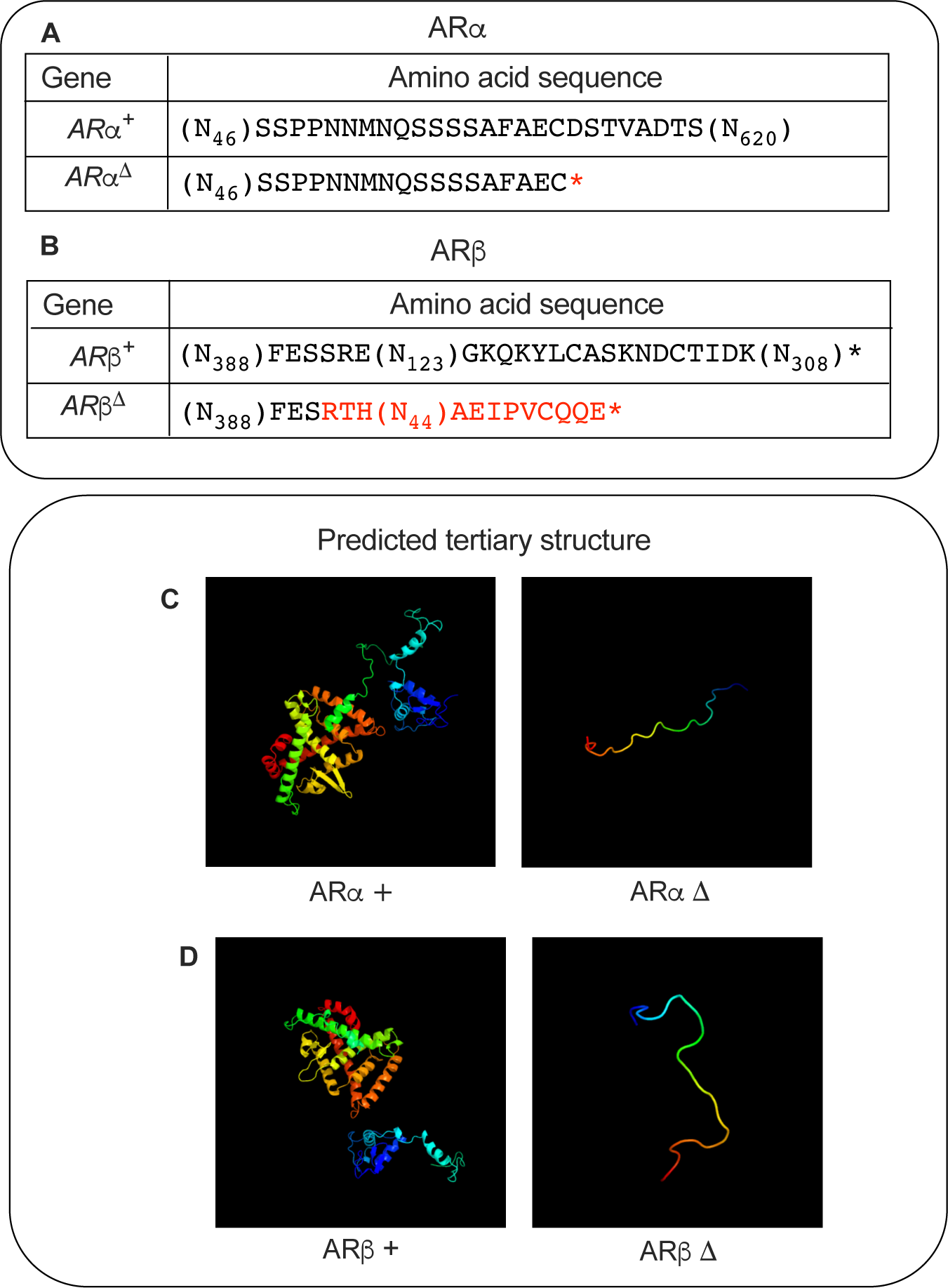
Predicted amino acid sequence and tertiary structure of wild-type and mutant forms of *AR*α and *ARβ*. **(A)** Amino acid sequence for the wild-type version of *AR*α (*AR*α^*+*^) and the mutant version of *AR*α (*AR*α^*Δ*^), with the latter containing a premature stop sequence (indicated by the red asterisk) induced by a frameshift mutation. **(B)** Amino acid sequence for the wild-type version of *ARβ* (*ARβ*^*+*^) and the mutant version of *ARβ* (*ARβ*^*Δ*^), with the latter containing incorrect amino acids (indicated in red font) and a premature stop sequence (indicated by the red asterisk) induced by a frameshift mutation. **(C)** Computer modeling (Phyre2(17)) predicts from the ARα^+^ amino acid sequence extensive tertiary structure in the ARα^+^ protein that is absent from that predicted from the ARα^Δ^ amino acid sequence. **(D)** Computer modeling (Phyre2(17)) predicts from the AR*β*^+^ amino acid sequence extensive tertiary structure in the ARα^+^ protein that is absent from that predicted from the AR*β*^Δ^ amino acid sequence. (N_#_) indicate the number of bases not shown in actual gene sequence for clarity. +=wild-type. *Δ*=frameshift deletion.

**Fig. S2.**
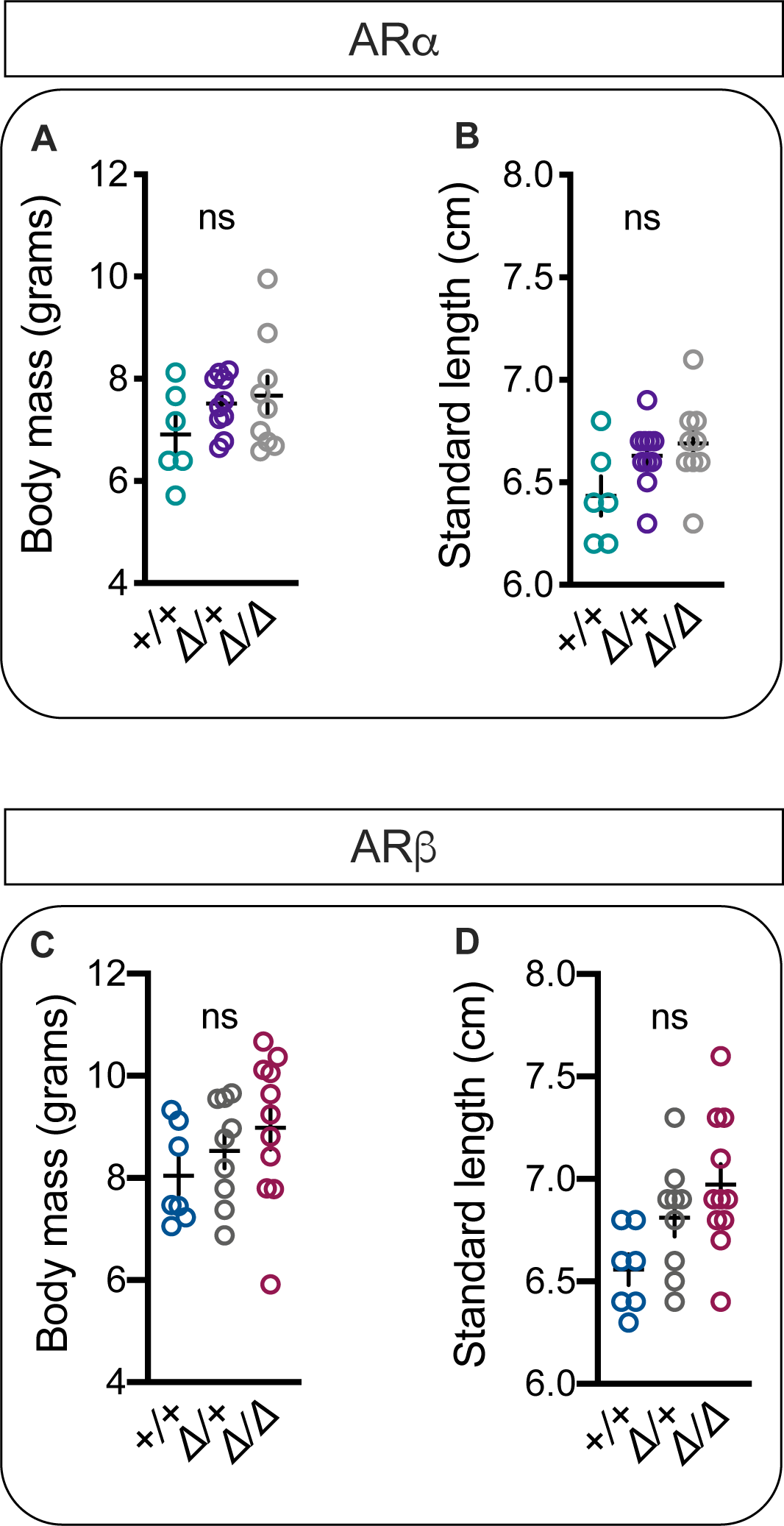
No effects of AR genotype on body mass and standard length. There was no effect of *ARα* (**A** and **B**) or *ARβ* (**C** and **D**) genotype on body size or standard length. +=wild-type. *Δ*=frameshift deletion. Circles represent data points for individual fish. Crosses represent Mean±SEM. ns=not significant.

**Fig. S3.**
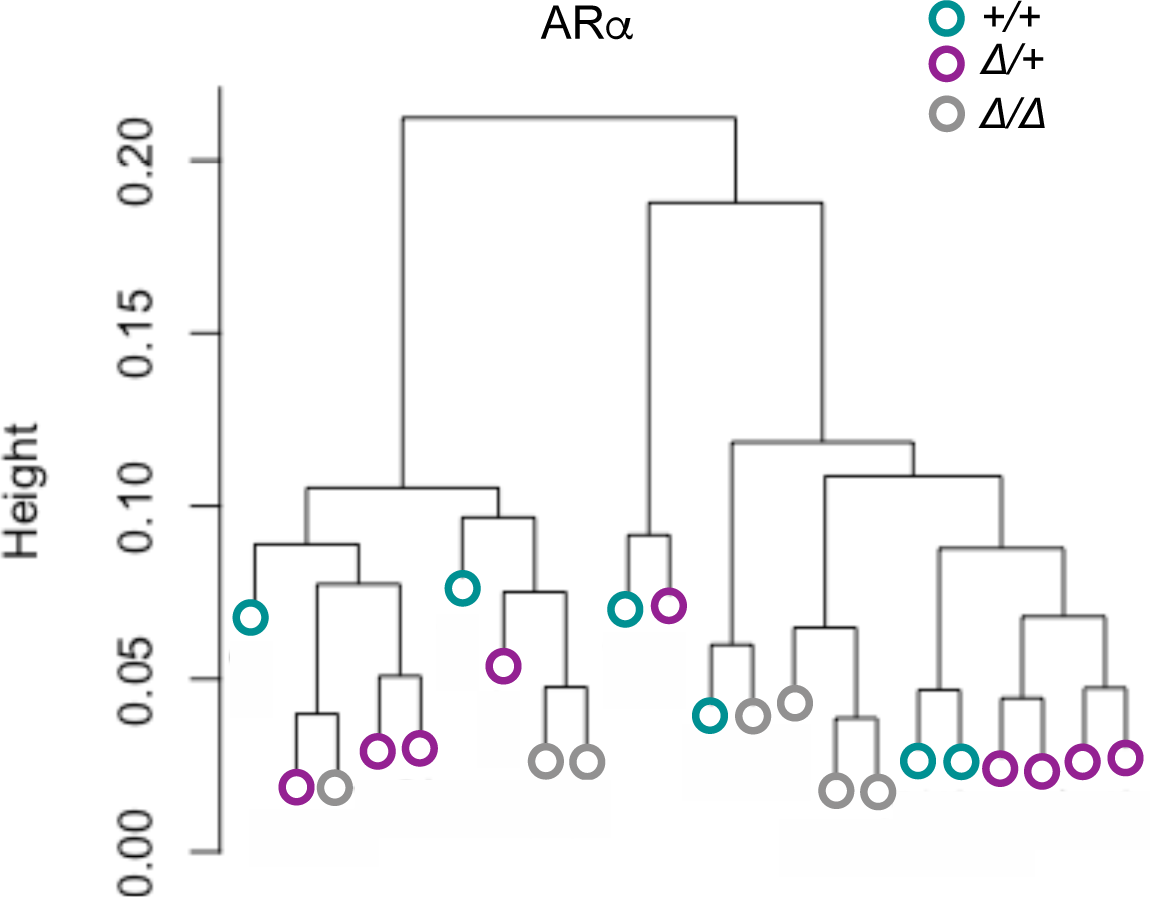
Results of hierarchical clustering of *ARα* fish by color. Each branch with a distinct colored circle at the tip represents a fish from a given genotype. Fish form clusters that are unrelated to genotype, confirming visual qualitative observations that *ARα* mutant males do not look different than *ARα*^*+/+*^ males.

**Fig. S4.**
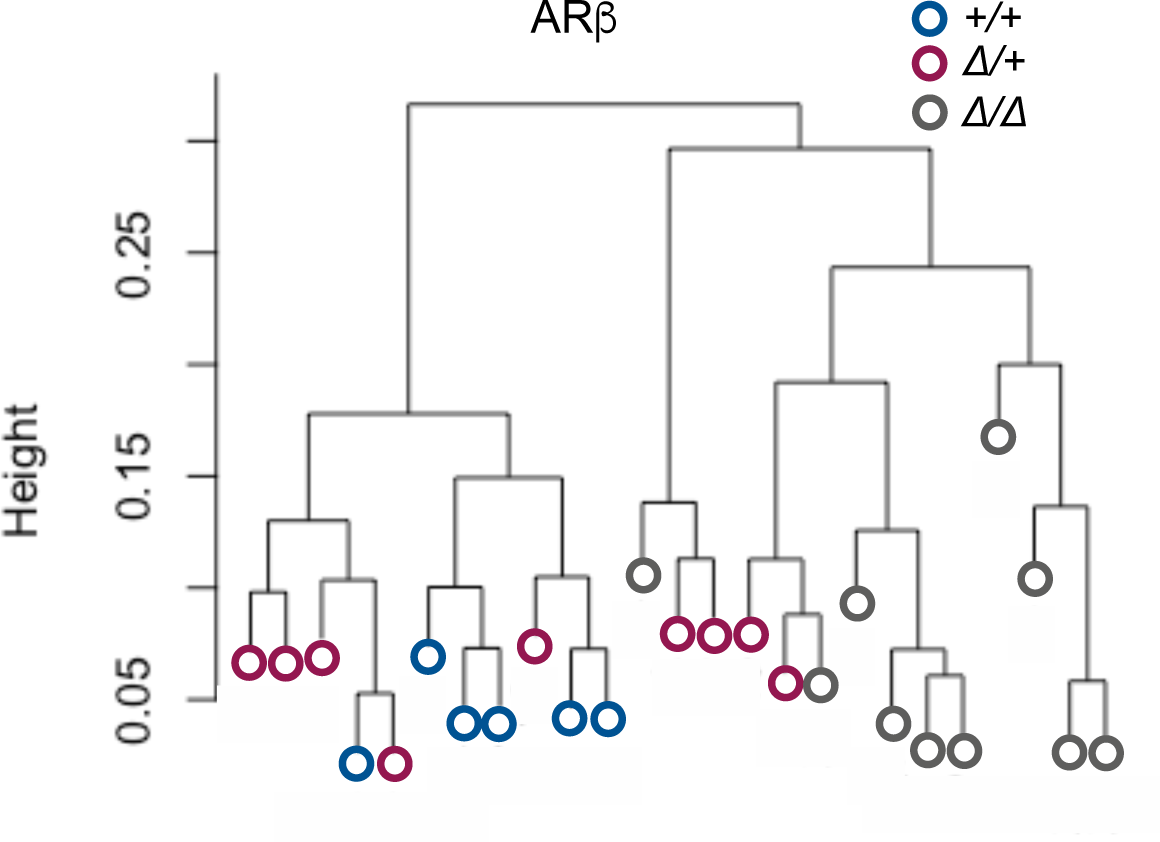
Results of hierarchical clustering of *ARβ* fish by color. Each branch with a distinct colored circle at the tip represents a fish from a given genotype. *ARβ*^*Δ/Δ*^and *ARβ*^*+/+*^ males fall into two distinct clusters, while *ARβ*^*Δ/+*^ males are split nearly equally between those two clusters, confirming visual qualitative observations that *ARβ*^*Δ/Δ*^ males look different than *ARβ*^*+/+*^ males and reflecting an intermediate coloration pattern in *ARβ*^*D/+*^ males.

**Fig. S5.**
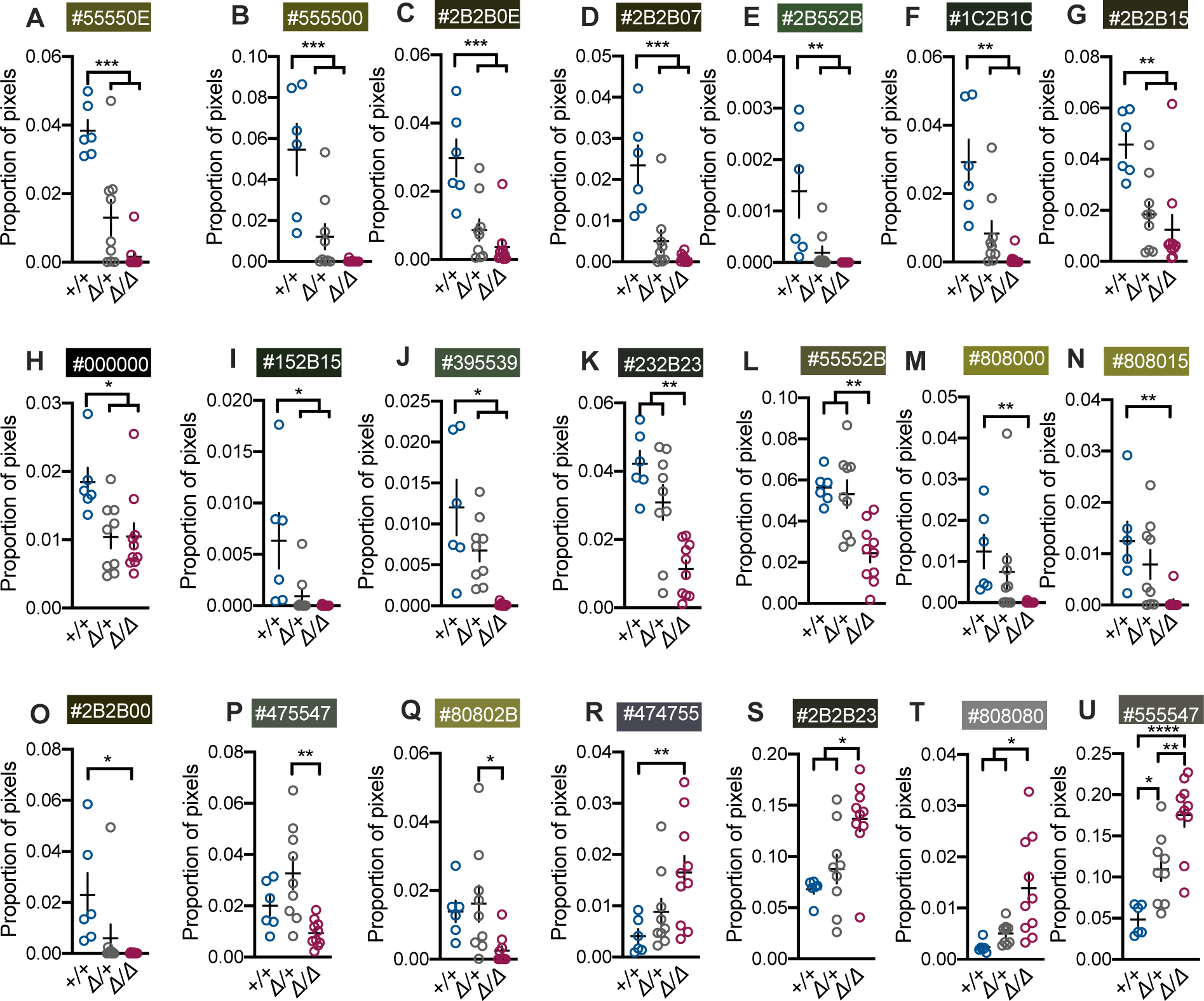
*ARβ* is necessary for the normal expression of numerous colors. (**A** to **J**) *ARβ*^*+/+*^ males exhibit a larger proportion of pixels for several colors compared to *ARβ* mutant males. (**K** and **L**) *ARβ* mutant males show a greater proportion of pixels for two colors compared to *ARβ*^*Δ/Δ*^ males. **(M** to **O**) *ARβ*^*+/+*^ males show a greater proportion of pixels for three colors compared to *ARβ*^*Δ/Δ*^ males but *ARβ*^*Δ/+*^ males were not different from either group for these colors. (**P** and **Q**) For two colors, *ARβ*^*Δ/+*^ males had a greater proportion of pixels than *ARβ*^*Δ/Δ*^ males but not *ARβ*^*+/+*^ males. **(R)** For one color, *ARβ*^*Δ/Δ*^ males had more pixels than *ARβ*^*+/+*^ males, **(S** to **T)** while for two colors *ARβ*^*Δ/Δ*^ males had a greater proportion of pixels compared to both groups. **(V)** *ARβ*^*Δ/+*^ males had more pixels for one color. +=wild-type. *Δ*=frameshift deletion. Circles represent data points for individual fish. Crosses represent Mean±SEM. \*\*\*\**P*<0.0001; \*\*\**P*<0.001; \*\**P* < 0.01; \**P* < 0.05.

**Fig. S6.**
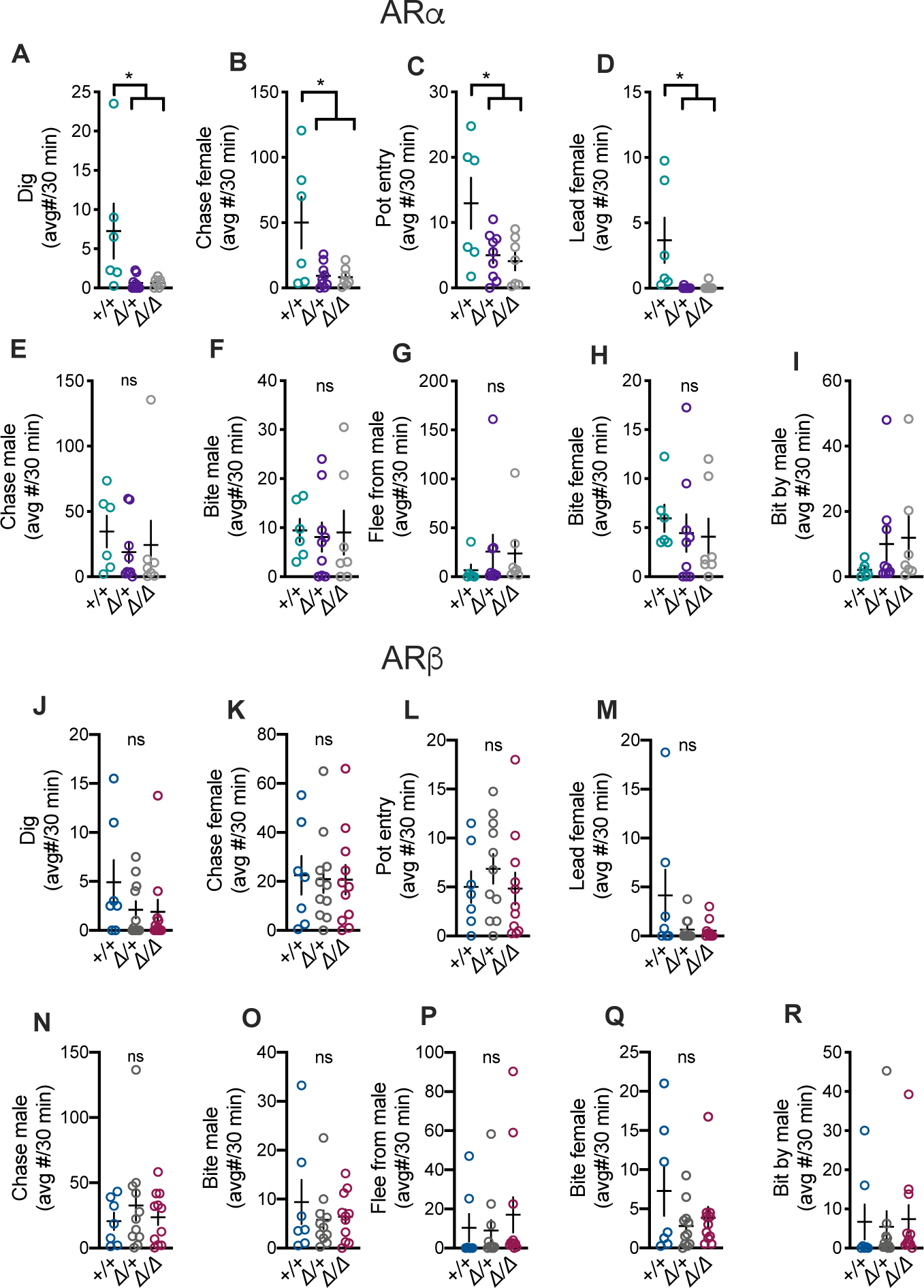
Effects of AR genotype on different social behaviors. (**A** to **D**) *AR*α is required for the performance of numerous reproductive behaviors. **(E** and **F)** *AR*α is not required for attacking the stimulus male. **(G)** *AR*α genotype does not affect the number of times males flee from the stimulus male. **(H and I)** *AR*α genotype does not affect the number of times males bite females or are bit by the stimulus male. (**J** to **M**) *ARβ* is not required for reproductive behaviors or (**N** and **O**) attacking the stimulus male. (**P**) *ARβ* genotype does not affect the number of times males flee from the stimulus male. (**Q** and **R**) *ARβ* genotype does not affect the number of times males bite females or are bit by the stimulus male. wt=wild-type. *Δ*=frameshift deletion. Circles represent data points for individual fish. Crosses represent Mean±SEM. ns=not significant. \**P* < 0.05.

**Fig. S7.**
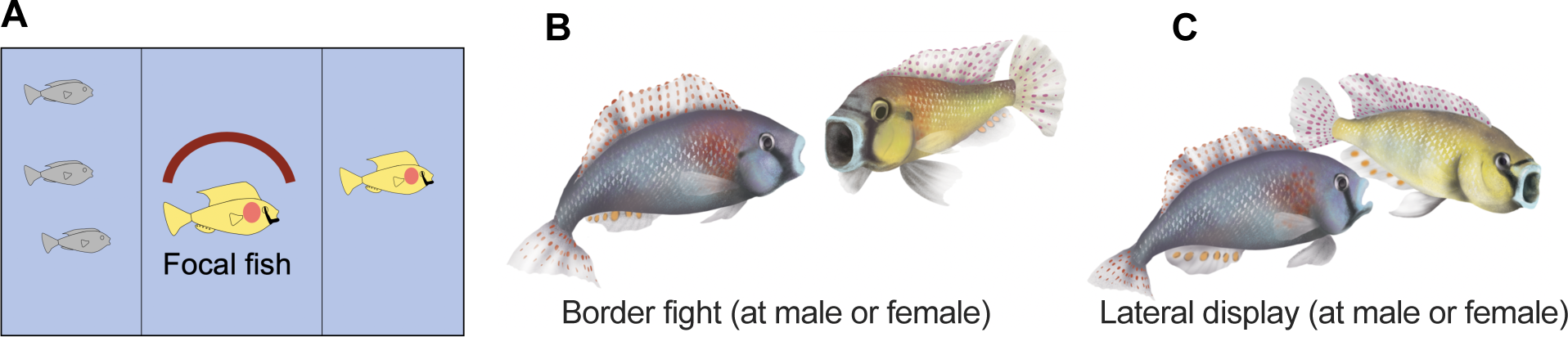
Scoring behavior during modified dyad assays. **(A)** A modified dyad assay was used to assess aggression. The focal fish was separated physically from the stimulus male and females by a transparent, perforated, acrylic barrier. (**B** and **C**) *A. burtoni* will perform border fights at either sex and perform lateral displays at either sex during modified dyad assays.

**Fig. S8.**
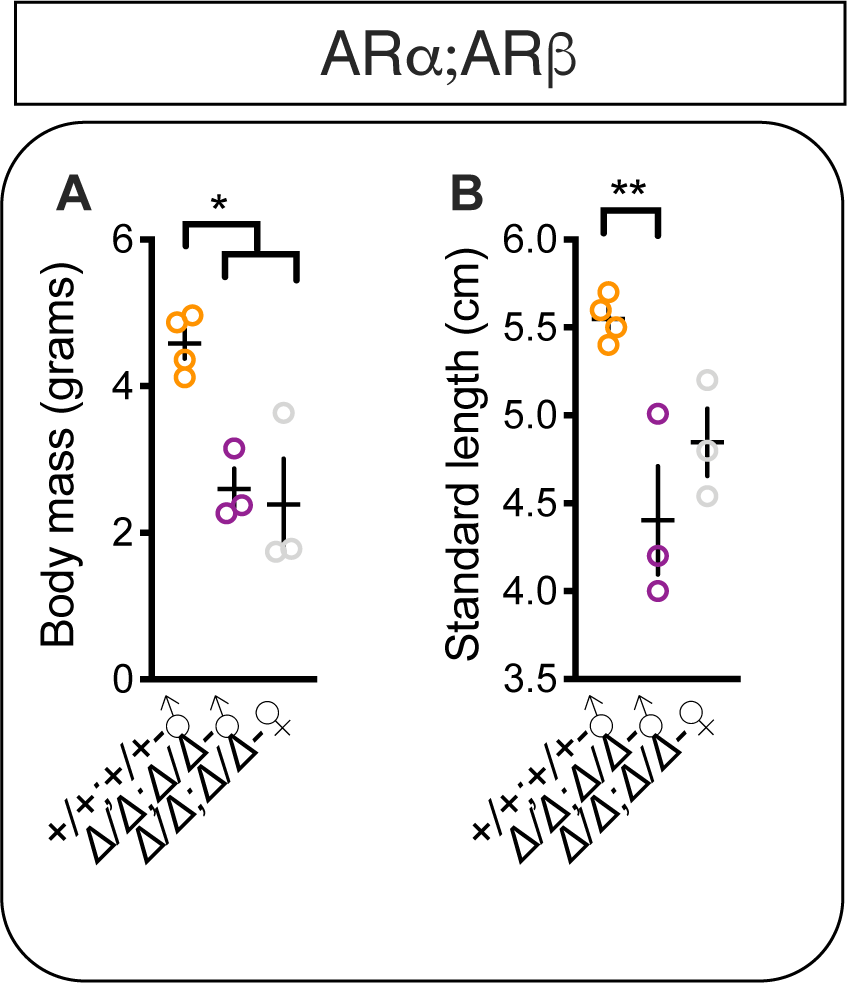
*ARα;ARβ* genotype affects body mass and standard length. **(A)** *ARα*^*+/+*^*;ARβ*^*+/+*^ males weighed more than *ARα*^*Δ/Δ*^*;ARβ*^*Δ/Δ*^ males and females **(B)** and were greater in standard length than *ARα*^*Δ/Δ*^*;ARβ*^*Δ/Δ*^ males. +=wild-type. *Δ*=frameshift deletion in both *ARα* and *ARβ* alleles. Circles represent data points for individual fish. Crosses represent Mean±SEM. \*\**P* < 0.01; \**P* < 0.05.

**Fig. S9.**
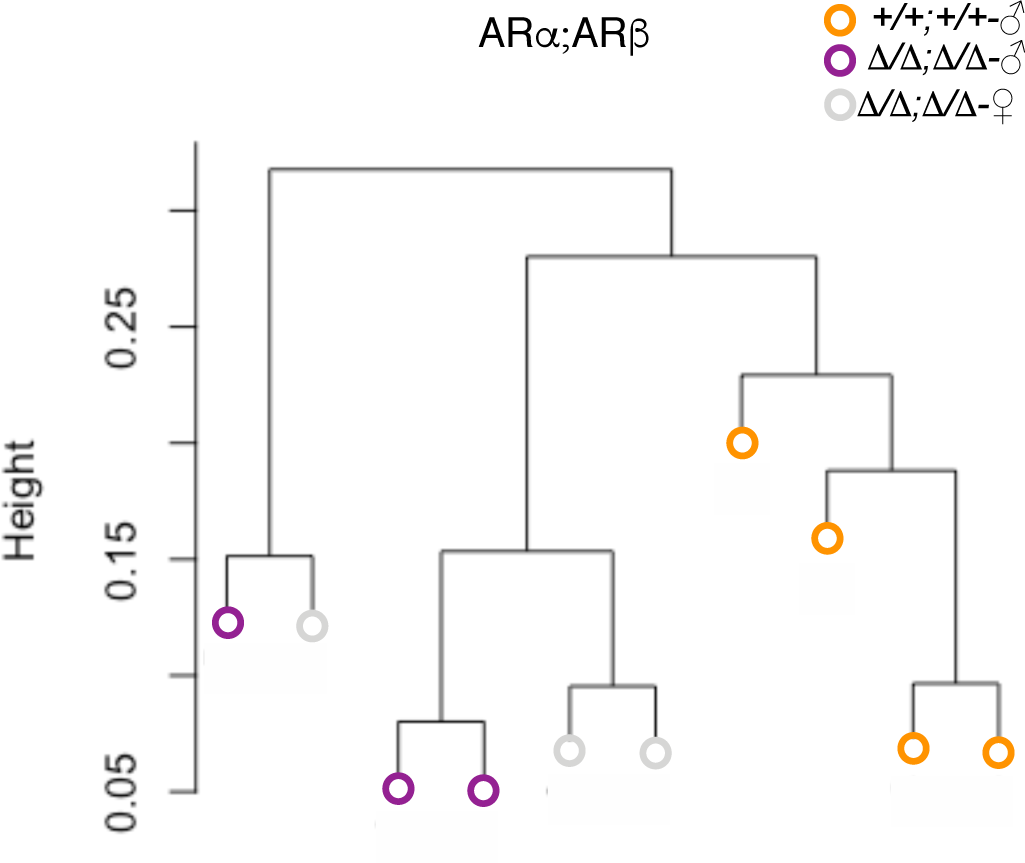
Results of hierarchical clustering of *ARα;ARβ* fish by color. Each branch with a distinct colored circle at the tip represents a fish from a given genotype. *ARα*^*+/+*^*;ARβ*^*+/+*^ males form a cluster that is distinct from *ARα*^*Δ/Δ*^*;ARβ*^*Δ/Δ*^ males and females, confirming visual qualitative observations that *ARα*^*+/+*^*;ARβ*^*+/+*^ males look different from *ARα*^*Δ/Δ*^*;ARβ*^*Δ/Δ*^ males and females *ARα*^*Δ/Δ*^*;ARβ*^*Δ/Δ*^.

**Fig. S10.**
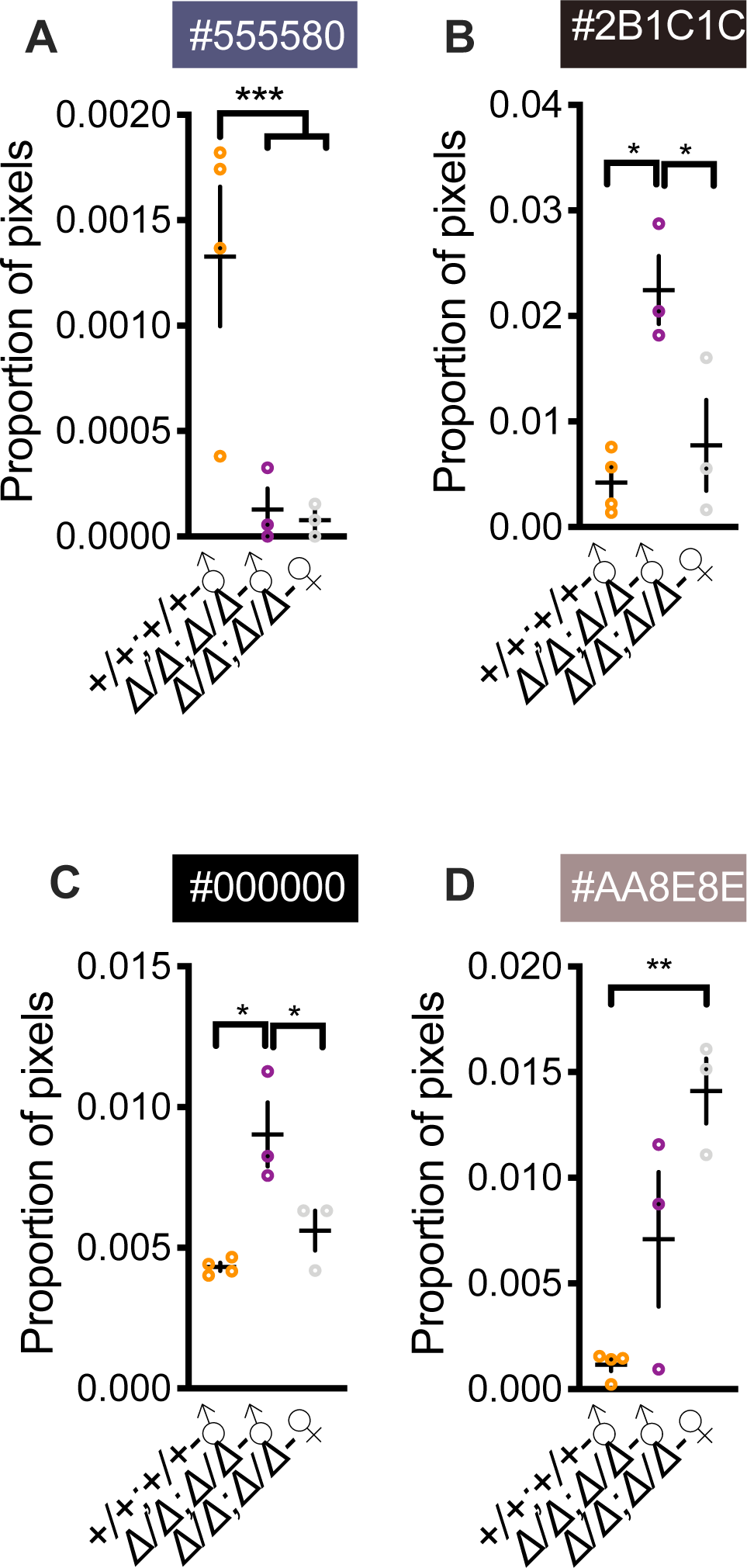
*ARα;ARβ* genotype affects whole-body coloration. **(A)** *ARα*^*+/+*^*;ARβ*^*+/+*^ males had a greater proportion of pixels for one color compared to than *ARα*^*Δ/Δ*^*;ARβ*^*Δ/Δ*^ males and females. **(B** and **C)** *ARα*^*Δ/Δ*^*;ARβ*^*Δ/Δ*^ males expressed more pixels for two colors than *ARα*^*+/+*^*;ARβ*^*+/+*^ males and *ARα*^*Δ/Δ*^*;ARβ*^*Δ/Δ*^ females. **(D)** *ARα*^*Δ/Δ*^*;ARβ*^*Δ/Δ*^ females expressed more pixels for one color compared to *ARα*^*+/+*^*;ARβ*^*+/+*^ males. +=wild-type. m=male. f=female. *Δ*=frameshift deletion in both *ARα* and *ARβ* alleles. Circles represent data points for individual fish. Crosses represent Mean±SEM. \*\*\**P*<0.001; \*\**P* < 0.01; \**P* < 0.05.

**Fig. S11.**
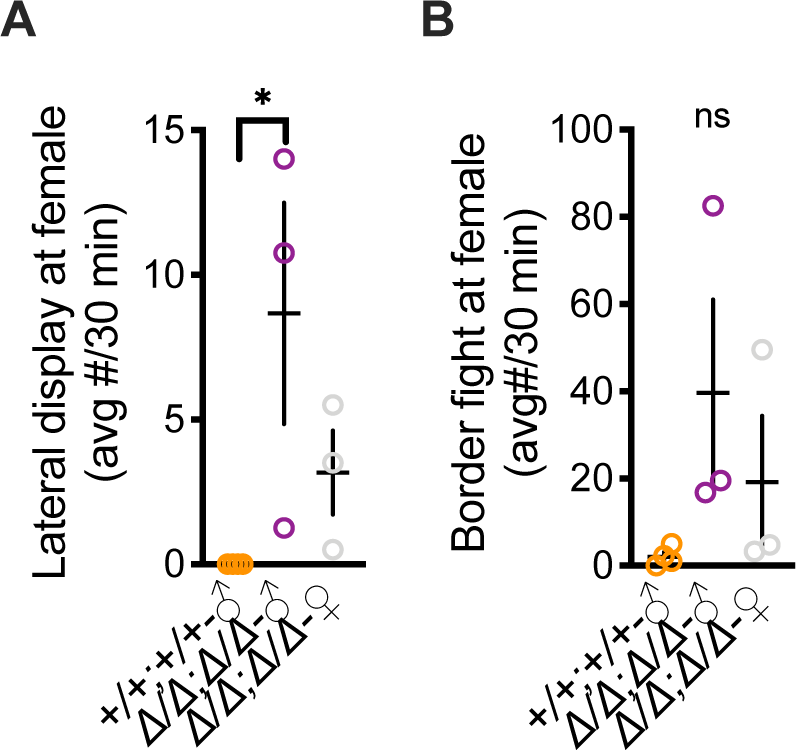
Effects on female-directed aggression. **(A)** *ARα*^*Δ/Δ*^*;ARβ*^*Δ/Δ*^ males performed more lateral displays directed towards females than *ARα*^*+/+*^*;ARβ*^*+/+*^ males. **(B)** There was a statistical trend (omnibus ANOVA *P*=0.06) for an effect of *ARα;ARβ* genotype on attacks directed towards females. +=wild-type. *Δ*=frameshift deletion. Circles represent data points for individual fish. Crosses represent Mean±SEM. \**P* < 0.05.

**Fig. S12.**
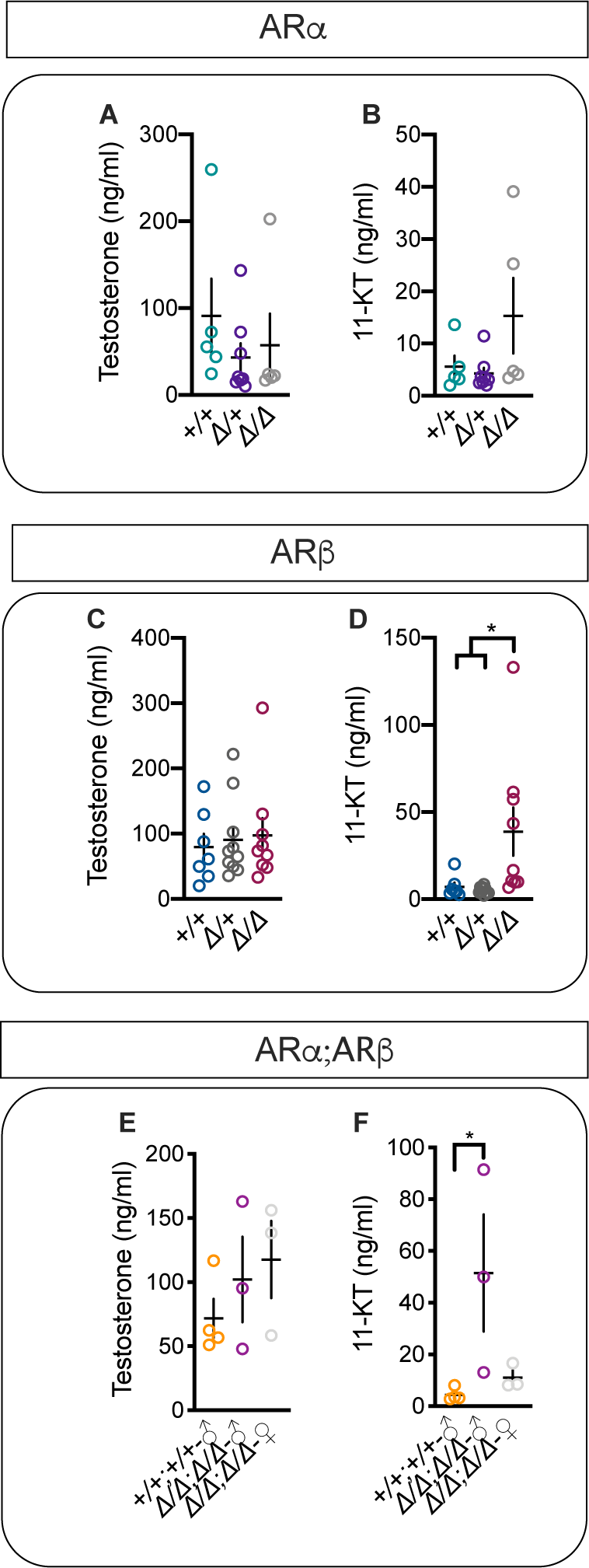
Effects on AR genotype on testosterone and 11-ketotestosterone levels. **(A** and **B)** *ARα* genotype has no effect on testosterone or 11-ketotestosterone (11-KT) levels. (**C**) *ARβ* genotype has no effect on testosterone levels, **(D)** but *ARβ*^*Δ/Δ*^ males have higher levels of 11-KT than *ARβ*^*Δ/+*^ and *ARβ*^*+/+*^ males. **(E)** *ARα;ARβ* genotype had no effect on testosterone levels, **(F)** but *ARα*^*Δ/Δ*^*;ARβ*^*Δ/Δ*^ had higher levels of 11-KT than *ARα*^*+/+*^*;ARβ*^*+/+*^ males. +=wild-type. *Δ*=frameshift deletion. Circles represent data points for individual fish. Crosses represent Mean±SEM. \**P* < 0.05.

### Supplementary Tables and Legends

**Table S1.** Gene information. Information was gathered from NCBI (https://www.ncbi.nlm.nih.gov/gene/?term=androgen+receptor+burtoni) on *A. burtoni* ARs.

| Gene Name/Alias | Description | Alias | Location |
| --- | --- | --- | --- |
| ar | androgen receptor | ARα | NW_005179415 |
| LOC102296288 | androgen receptor-like | ARβ | NW_005179497 |

**Table S2.**
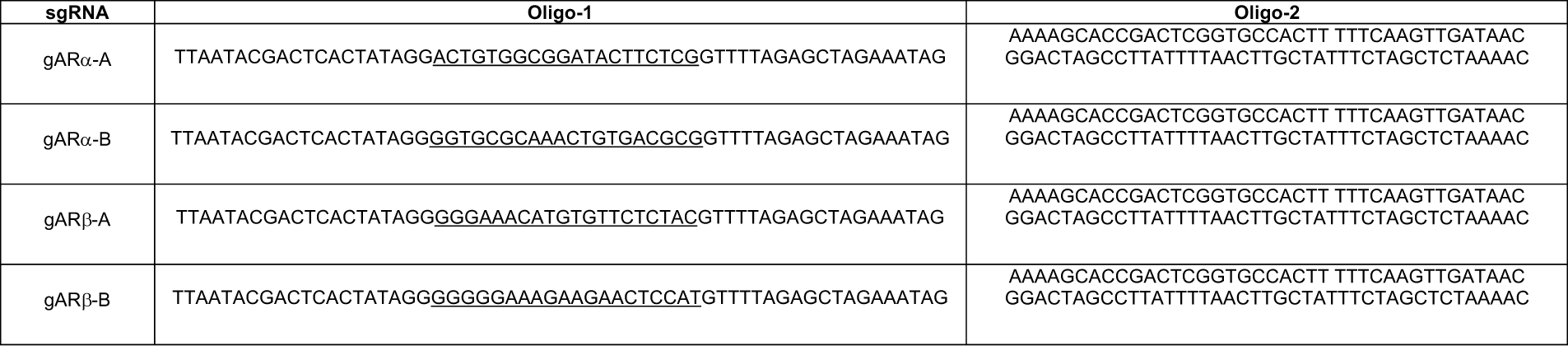
Oligonucleotides used to synthesize single-guide RNA (sgRNA). We synthesized sgRNA by annealing oligo-1, which was specific to each target site, to oligo-2, a generic oligo that allows for amplification of the sgRNAs. g=guide. AR=androgen receptor.

## Notes

### Competing Interest Statement

The authors have declared no competing interest.

## References

1. R. M. Sapolsky, The influence of social hierarchy on primate health. 308, 648–653 (2005).

2. R. D. Fernald, Cognitive skills needed for social hierarchies. Cold Spring Harb. Symp. Quant. Biol. 79, 229–236 (2014).

3. R. D. Fernald, K. P. Maruska, Social information changes the brain. Proc. Natl. Acad. Sci. 109, 17194–17199 (2012).

4. J. Dreher, S. Dunne, A. Pazderska, T. Frodl, J. J. Nolan, Testosterone causes both prosocial and antisocial status-enhancing behaviors in human males. Proc. Natl. Acad. Sci. 113, 11633–11638 (2016).

5. B. A. Alward, A. T. Hilliard, R. A. York, R. D. Fernald, Hormonal regulation of social ascent and temporal patterns of behavior in an African cichlid. Horm. Behav. 107, 83–95 (2019).

6. C. Eisenegger, J. Haushofer, E. Fehr, The role of testosterone in social interaction. Trends Cogn. Sci. 15, 263–271 (2011).

7. Z. A. Knight, K. M. Shokat, Chemical Genetics: Where Genetics and Pharmacology Meet. Cell 128, 425–430 (2007).

8. T. J. Stevenson, et al., The value of comparative animal research: Krogh’s Principle facilitates scientific discoveries. Policy Insights from Behav. Brain Sci. 5, 118–125 (2018).

9. G. E. Robinson, R. D. Fernald, D. F. Clayton, Genes and social behavior. Science 322, 896–900 (2008).

10. L. A. O’Connell, H. A. Hofmann, Social status predicts how sex steroid receptors regulate complex behavior across levels of biological organization. Endocrinology 153, 1341–1351 (2012).

11. D. Brawand, et al., The genomic substrate for adaptive radiation in African cichlid fish. Nature 513, 375–381 (2014).

12. S. Ohno, Evolution by gene duplication (Springer, 1970).

13. J. A. Doudna, E. Charpentier, The new frontier of genome engineering with CRISPR-Cas9. Science 346, 1–9 (2014).

14. L. E. Jao, S. R. Wente, W. Chen, Efficient multiplex biallelic zebrafish genome editing using a CRISPR nuclease system. Proc. Natl. Acad. Sci. U. S. A. 110, 13904–13909 (2013).

15. G. K. Varshney, et al., High-throughput gene targeting and phenotyping in zebrafish using CRISPR/Cas9. Genome Res. 25, 1030–1042 (2015).

16. S. A. Juntti, et al., A neural basis for control of cichlid female reproductive behavior by prostaglandin F2α. Curr. Biol. 26, 943–949 (2016).

17. L. A. Kelley, S. Mezulis, C.M. Yates, M.N Wass, & M.J. Sternberg, The Phyre2 web portal for protein modeling, prediction and analysis. Nat. Protoc. 10, 845–858 (2015).

18. K. P. Maruska, R. D. Fernald, Behavioral and physiological plasticity: Rapid changes during social ascent in an African cichlid fish. Horm. Behav. 58, 230–240 (2010).

19. S. S. Burmeister, E. D. Jarvis, R. D. Fernald, Rapid behavioral and genomic responses to social opportunity. PLoS One 3, 1–9 (2005).

20. R. M. Alcazar, L. Becker, A. T. Hilliard, K. R. Kent, R. D. Fernald, Two types of dominant male cichlid fish: behavioral and hormonal characteristics. Biol. Open 5, 1061–1071 (2016).

21. S. C. P. Renn, E. J. Fraser, N. Aubin-Horth, B. C. Trainor, H. A. Hofmann, Females of an African cichlid fish display male-typical social dominance behavior and elevated androgens in the absence of males. Horm. Behav. 61, 496–503 (2012).

22. S. A. Juntti, et al., The androgen receptor governs the execution, but not programming, of male sexual and territorial behaviors. Neuron 66, 260–72 (2010).

23. T. Sato, et al., Brain masculinization requires androgen receptor function. Proc. Natl. Acad. Sci. U. S. A. 101, 1673–1678 (2004).

24. X. B. Cheng, et al., Characterizing the neuroendocrine and ovarian defects of androgen receptor-knockout female mice. Am. J. Physiol. - Endocrinol. Metab. 305 (2013).

25. J. K. Desjardins, H. A. Hofmann, R. D. Fernald, Social context influences aggressive and courtship behavior in a cichlid Fish. PLoS One 7, 1–5 (2012).

26. C. C. Chen, R. D. Fernald, Visual information alone changes behavior and physiology during social interactions in a cichlid fish (Astatotilapia burtoni). PLoS One 6, 1–12 (2011).

27. B. A. Alward, C. A. Cornil, J. Balthazart, G. F. Ball, The regulation of birdsong by testosterone: Multiple time-scales and multiple sites of action. Horm. Behav. 104, 32–40 (2018).

28. J. M. Carré, J. Archer, Testosterone and human behavior: the role of individual and contextual variables. Curr. Opin. Psychol. 19, 149–153 (2018).

29. C. M. Crowder, C. S. Lassiter, D. A. Gorelick, Nuclear Androgen Receptor Regulates Testes Organization and Oocyte Maturation in Zebrafish. Endocrinology 159, 980–993 (2018).

30. L. Yong, Z. Thet, Y. Zhu, Genetic editing of the androgen receptor contributes to impaired male courtship behavior in zebrafish. J. Exp. Biol. 220, 3017–3021 (2017).

31. M.E. Santos, W. Salzburger, How cichlids diversify. Science 338, 619–620 (2012).

32. R. D. Fernald, N. R. Hirata, Field study of Haplochromis burtoni: Quantitative behavioural observations. Anim. Behav. 25, 964–975 (1977).

33. G. K. Varshney, et al., A high-throughput functional genomics workflow based on CRISPR/Cas9-mediated targeted mutagenesis in zebrafish. Nat. Protoc. 11, 2357–2375 (2016).

34. H. E. Fox, S. A. White, M. H. Kao, R. D. Fernald, Stress and dominance in a social fish. J. Neurosci. 17, 6463–6469 (1997).

